# A decision-support framework to optimize border control for global outbreak mitigation

**DOI:** 10.1101/414185

**Authors:** Aleksa Zlojutro, David Rey, Lauren Gardner

**Affiliations:** School of Civil and Environmental Engineering, University of New South Wales (UNSW) Sydney, Sydney, NSW, 2052, Australia; Department of Civil Engineering, Johns Hopkins University, Baltimore, MD 21218, USA

## Abstract

The introduction and spread of emerging infectious diseases is increasing in both prevalence and scale. Whether naturally, accidentally or maliciously introduced, the substantial uncertainty surrounding the emergence of novel viruses, specifically where they may come from and how they will spread, demands robust and quantifiably validated outbreak control policies that can be implemented in real time. This work presents a novel mathematical modeling framework that integrates both outbreak dynamics and outbreak control into a decision support tool for mitigating infectious disease pandemics that spread through passenger air travel. An ensemble of border control strategies that exploit properties of the air traffic network structure and expected outbreak behavior are proposed. A stochastic metapopulation epidemic model is developed to evaluate and rank the control strategies based on their effectiveness in reducing the spread of outbreaks. Sensitivity analyses are conducted to illustrate the robustness of the proposed control strategies across a range of outbreak scenarios, and a case study is presented for the 2009 H1N1 influenza pandemic. This study highlights the importance of strategically allocating outbreak control resources, and the results can be used to identify the most robust border control policy that can be implemented in the early stages of an outbreak.

## Introduction

The potential harm posed by the introduction and spread of emerging infectious diseases has been recently illustrated by the 2009 H1N1 ^1^, SARS ^2^, and Zika ^3^ epidemics. There is considerable evidence that such pandemics are likely to become more frequent unless action is taken to mitigate their spread at a global scale ^4-7^. For this reason, resilience management for outbreaks has attracted a growing body of literature, both from epidemic modelers and the optimization and control community. One of the most critical aspects of resilience management is the need to combine accurate epidemic growth models with detailed outbreak control strategies ^8^, representing a gap in the literature which this study aims to fill.

The availability of large scale data and growing computational capabilities has significantly advanced infectious disease spread models in recent years ^9-11^. The community has notably explored the impact of network topology, epidemic thresholds and diffusion models on outbreak spread patterns ^12-15^. A range of epidemic models have been developed, which increase in complexity from single-population, deterministic models to metapopulation, stochastic models ^6,16,17^. Deterministic models provide efficient mathematical representation but lack the realism of stochastic simulation models ^17-20^. On the other hand, detailed computational and visualization tools such as GLEaM ^21^ and STEM ^22^, among others ^23-26^, have emerged as powerful solutions to model the spread of infectious diseases with a high level of accuracy and even measure the impact of control strategies ^27^, albeit at a high computational cost. As an alternative to simulation-based models, analytical global epidemic models have also been developed to characterize the stochastic spread of infectious diseases in metapopulation networks ^5,28,29^.

As highlighted by ^4^, the intensive restriction of human activities in metapopulation networks may undermine the system’s functionality and lead to significant societal costs. Hence, in efforts to mitigate large-scale pandemics, it is critical to cautiously deploy control strategies to minimize disruptions and maximize the reactiveness of the system ^30-32^. At a global scale, passenger air travel is known to play a critical role in the spread of infectious disease ^15,23,33-35^. Additionally, border control has been shown to play a pivotal role in mitigating epidemics, especially during the emerging stage of outbreaks ^5,36-39^. Border control is typically deployed at airports in an attempt to prevent the spread of an infectious disease between cities, states and countries through passenger air travel ^40^. However, identifying the optimal set of airports for deploying border control is challenging, especially when only limited control resources are available, *e.g.*, budget constraints.

In this work we present a novel mathematical modeling framework that integrates both outbreak dynamics and outbreak control into a decision-support tool for mitigating infectious disease pandemics at the onset of an outbreak through border control. The border control mechanism considered in this work is passenger screening upon arrival at airports (entry screening), which is used to identify infected or at-risk individuals and provide immediate treatment and isolation to reduce the risk of further transmission ^41^. The uncertainty of exit screening effectiveness in other countries combined with the possible development of symptoms during a flight ^42^ has prompted several governments to deem entry screening as crucial to the protection of their countries ^43^, and further motivates its use in this study. The proposed model seeks to determine the optimal set of airports that should be allocated screening resources (technology and personnel), and the corresponding amount (thus dictating the proportion of arriving passengers that can be screened) such that the outbreak risk is minimized. We propose an ensemble of control strategies that exploit the heterogeneity of the air traffic network structure and outputs from an outbreak simulation model in the allocation of control resources. We evaluate each control strategy using a stochastic metapopulation epidemic model, and compare the strategies based on their effectiveness in reducing and the spread of outbreaks. Note that we are not proposing or evaluating air travel restrictions, which have substantial economic costs as well as recognized limitations in their ability to prevent or reduce the scale of pandemics ^20,29,44-48^. The goal of the proposed decision-support framework is to optimize the use of airport screening as a method of border control to delay the introduction of a new disease into susceptible cities, thus providing local public health authorities more time to plan, prepare and distribute local control strategies, *e.g.*, anti-virals, vaccines, source exit screening etc., which must be rapidly administered if/when infection is introduced.

This study builds upon previous work ^49,50^ that presented a mathematical modeling framework to integrate control and outbreak dynamics. We extend this line of work through the following substantial contributions: 1) we model outbreak dynamics using a stochastic metapopulation framework as opposed to a deterministic model, 2) we propose and evaluate a novel set of control strategies, 3) the control strategies are embedded within a resource constrained decision-support framework, 4) the model is calibrated using historical outbreak data, and 5) a case study is presented for the 2009 H1N1 outbreak. Additionally, a range sensitivity analysis is conducted to illustrate the robustness of the model with regards to variability across outbreak scenarios, disease parameters, policy considerations, and modelling assumptions. The analysis elucidates key trade-offs in terms of budget availability and outbreak mitigation which can be leveraged to inform on public health policy for global epidemic preparedness and control.

## Materials and Methods

### Mathematical Model

In this section, we present both the stochastic metapopulation epidemic model used to simulate outbreak dynamics, and its integration within the proposed decision-support framework. Our metapopulation model is based on a global air travel network which connects local, city-level, populations. Formally, the proposed metapopulation network can be represented by a graph *G* = (*V, E*) where *V* is the set of nodes and *E* is the set of directed edges in the network. Nodes represent cities and edges represent passenger travel routes, possibly including stopovers, among cities. At each node of the network, we locally model outbreak dynamics using a discrete-time Susceptible-Exposed-Infected-Recovered (SEIR) compartmental model ^51^. The time steps are set to be *t* ∈ *T* = {1, 2, …, *t*_*obs*_} where *t*_*obs*_ is the time step where the state of the outbreak is being evaluated. Local and global outbreak dynamics models are coupled by indexing compartmental states by network nodes *i* ∈ *V* and time steps *t*. Specifically, we denote *S*_*i,t*_, *E*_*i,t*_, *I*_*i,t*_ and *R*_*i,t*_ the susceptible, exposed, infectious and recovered compartments at node *i* at time *t*. Because our objective is to prevent spread in the early stages of an outbreak (*e.g.,* weeks or months), we assume that nodes have time-independent populations and we denote *N*_*i*_ the population at node *i* ∈*V*. We aim to use this metapopulation model to capture day-to-day global travel dynamics, hence time steps are assumed to be of the order of magnitude of a day in length.

We use a multi-commodity network flow model with time-dependent edge flows to model passenger movements from their origin node to their destination node. Let Π_*i,j*_ be the set of paths from *i* ∈ *V* to *ij,tj* ∈ *V*. We denote 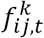 the average passenger flow on the route from *i* ∈ *V* to *j* ∈ *V* using path *k* ∈ Π_*ij*_ at time step *t* and we assume symmetric passenger flows for all pairs of origins and destinations, i.e. 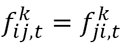. We denote 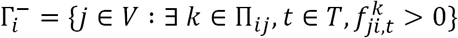and 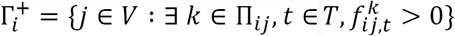 the sets of nodes connected to and from node *i* ∈ *V*, respectively. Each path *k* ∈ Π_*ij*_ is an ordered sequence of nodes starting at *i* and ending at *j*, i.e. *k* = ⟨*i, n*_1_, *n*_2_, …, *j*⟩ and we denote *n*_1_ ≺ *n*_2_ the precedence relationship among nodes in the path.

The governing infection dynamics of the SEIR model ^51^ are used to model local outbreak dynamics in each city. For the purposes of this work the contact rate is assumed to be constant across populations. We denote *β*_*i*_ the (local) contact rate at node *i, γ* the transition or recovery rate and α the exposed parameter. In addition, we define *λ* ∈ [0,1] the likelihood to travel when infectious, with *λ* = 1 representing the case where infected and healthy individuals are equally likely to travel. This parameter aims to represent the impact of reduced travel demand when infectious individuals are unable to travel due to severe symptoms. Finally, we assume that compartmental edge flows are proportional to tail node states, i.e. the number of travelers in a state is proportional to the number of individuals in this state at the origin node. Discrete-time stochastic metapopulation outbreak dynamics are summarized in Equation (1) below.

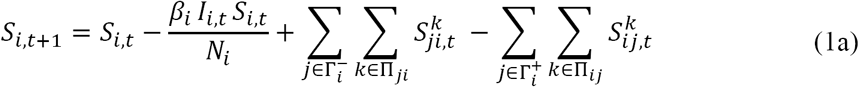

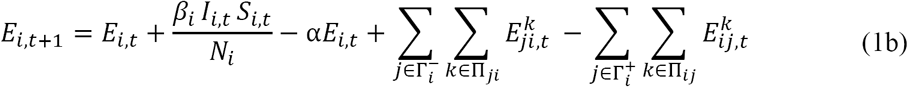

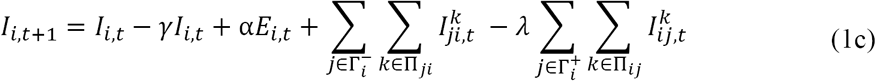

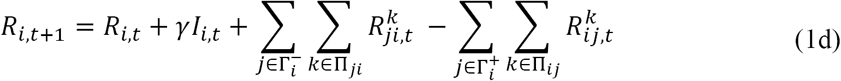

The symbols 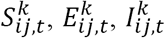 and 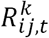 represent compartmental edge flows on (*i, j*) with destination *k* at time step *t*. For compartments *S* and *R*, compartmental edge flows are assumed deterministic and equal to their expected values, i.e.: 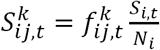 and 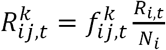. However, since the compartmental edge flows of exposed and infectious passengers may be considerably smaller than that of other compartments, we model 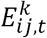 and 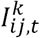 as discrete random variables, as the stochastic allocation of infected individuals to destinations is critical when modeling the early stages of an outbreak. Specifically, we define these compartmental edge flows as follows:

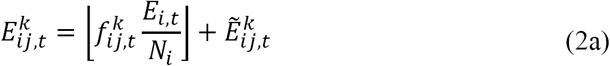

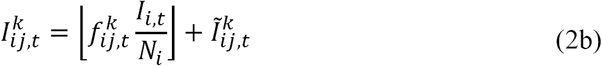

where 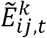 and 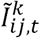 are discrete random variables representing the number of exposed and infectious passengers, respectively, beyond the integer-part of their respective compartmental edge flows. Further,let 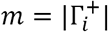 be the number of destination nodes from node *i* ∈ *V*, the vector 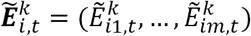 (resp 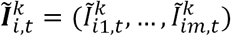 follows a multinomial distribution with a number of trials 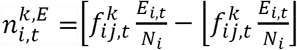 (resp. 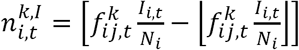) and probability vector 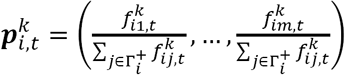

This stochastic formulation aims to model integer, compartmental edge flows and prevent the movement of fractional exposed or infectious individuals which may result in unrealistic epidemic behavior at a global scale ^52^. Consequently, the compartmental states *S*_*i,t*_,*E* _*i,t*_, *I*_*i,t*_ and *R*_*i,t*_ are also random variables representative of the evolution of the outbreak over time and space.

To integrate control decisions within the above stochastic metapopulation network we model passenger screening upon arrival at airports as a control variable. Passenger screening can be done through visual inspections of passengers, health declaration cards and/or infrared thermal image scanners ^53^. In this work the specific type of screening is not the focus; as the framework is applicable to multiple control mechanisms. We propose to use airport screening rates as the main control variables, which are representative of the proportion of arriving passengers successfully screened at a given airport. We denote *x*_*i,t*_ ∈ [0,1] the control rate at node *i* at time step *t*. Control variables can be incorporated in the proposed metapopulation epidemic model by re-defining Equations (1c) and (1d) as follows:

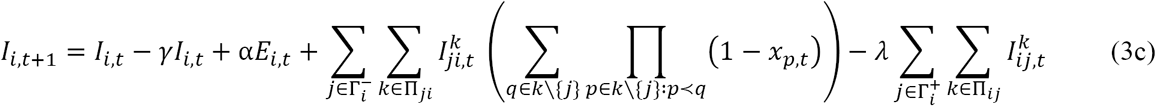

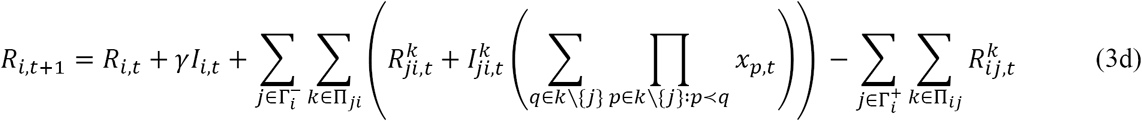

This formulation is able to capture the combined effects of screening passengers at multiple nodes along their travel route. The combination of Equations (1a), (1b), (3c) and (3d), hereby to as (3), can be viewed as a control-driven stochastic metapopulation epidemic model wherein variables *x*_*i,t*_ represent the level of control over time space in the network. A control rate of less than one can be interpreted as shortcomings of the methods or technology involved with screening. We assume that passengers coming into a controlled airport who are successfully identified as infected individuals are isolated for treatment, and hence are no longer able to spread infection. In terms of the model, infected individuals screened at a controlled node are assumed to transition to the recovered state *R*. The main objective function is to minimize the expected cumulative number of infected individuals at the observation time ^27^. This can be stated as follows:

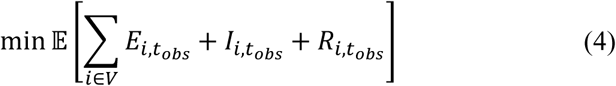

The challenge in policy decision making results because of a constraint on available resources. To address this challenge we introduce a budget and cost for control. The cost of using outbreak control resources is modeled using a generic cost function *c*_*i*_ which maps control variables to monetary costs. We assume that setup costs *s*_*i*_, are associated with the deployment of control resources at a node *i* ∈ *V*, e.g., installation of new screening technologies and training of personnel. To model the activation of control at a node, we introduce binary variables *y*_*i*_ ∈ {0,1} which take value 1 if node *i* is allocated a non-zero amount of control resources. We assume that the cost of deploying control resources at *i* over the control period *T* is a function of the total incoming edge flow to *i* at each time step *t*, i.e. 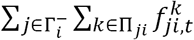 and we denote 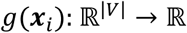 the variable part of the cost function, where ***x***_*i*_ ∈ [0,1]^|*T*|^ is the local control vector at node *i* ∈ *V*. The variable portion of the cost function is representative of a per passenger screening cost each day. A generic form of the cost function *c*_*i*_ is then:

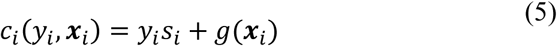

In addition, control and setup variables are linked through the inequalities *x*_*i,t*_ ≤ *y*_*i*_ indicating that setup costs are incurred if control resources are deployed at any time during the control period. Finally, we assume that a budget *B* is available to for deploying control resources which translates into the budget constraint:

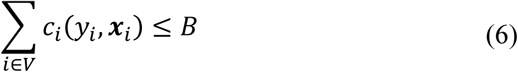

Our objective is to optimize the control vector ***x*** ∈ [0,1]^|*V*||*T*|^ such that the impact of the outbreak at *t*_*obs*_ as represented by (4) is minimized subject to control resources constraints and outbreak dynamics as governed by the stochastic metapopulation epidemic model (3) wherein compartmental edge flows *E*_*ij,t*_ and *I*_*ij,t*_ as well as compartmental states *S*_*i,t*_, *E*_*i,t*_, *I*_*i,t*_ and *R*_*i,t*_ are discrete random variables. This optimization formulation is summarized in Eq (7) below.

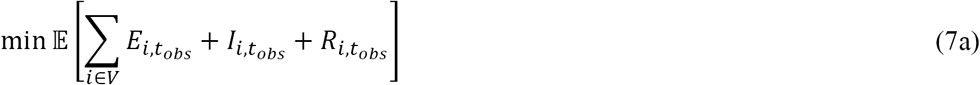

Subject to:

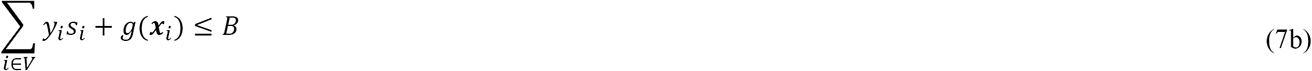

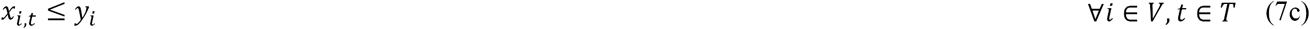

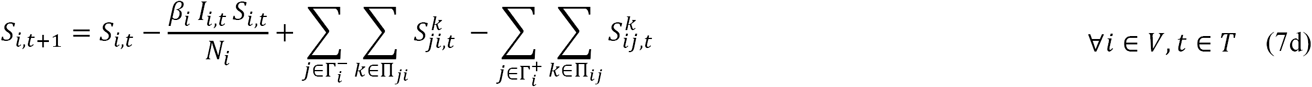

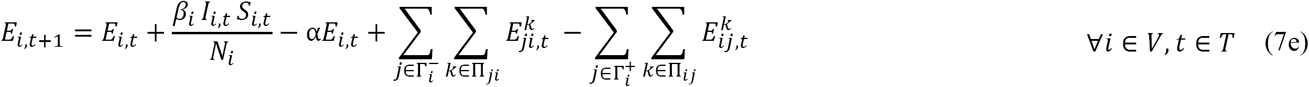

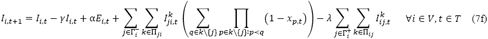

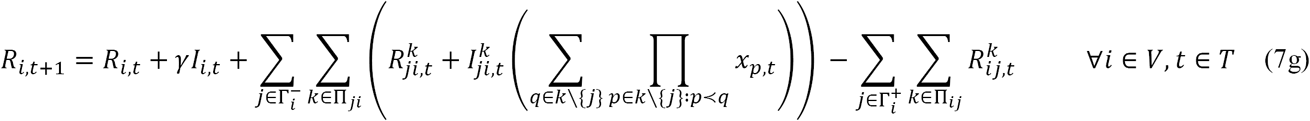

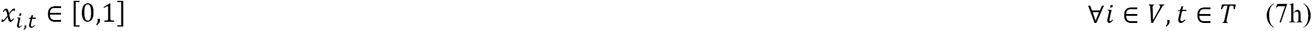

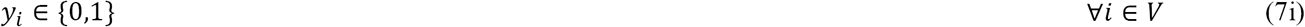

### Proposed Control Strategies

The final outbreak dynamics model with control decisions incorporated can be used as a tool to solve the resource allocation problem, and evaluate various control strategies. We propose a set of control strategies to optimize the control vector ***x*** subject to resource constraints. In this work, each proposed control strategy relies on a different metric to rank airports, and this ranking is then used to allocate control resources to a select set of airports. The objective of all strategies is to minimize the impact of the outbreak at a pre-selected future date we call the observation time, *t*_*obs*_, at which impact is measured both in terms of total cumulative cases and number of infected cities.

Although the proposed model can accommodate dynamic control strategies, for the purposes of this work we focus on static control strategies, wherein nodes are controlled at the same level throughout the period of observation. Hence, we define and use the following static-equivalent cost function *g*_s_ (*x*_*i*_): ℝ → ℝ which we assume to be invertible over its domain. Our approach is based on a greedy resource allocation algorithm which iterates over a sorted set of nodes 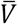 until the outbreak control budget is depleted. The pseudo-code of this resource allocation procedure is summarized in Algorithm 1.

#### Algorithm 1: Greedy outbreak control resource allocation

**1 Input:** Sorted node set 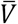 budget *B*

**2 Output:** Control vector ***x***

3 ***x* ← 0**

4 *U* ← 0

5 **for** *i* in 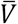

6 **if** *U* + *s*_*i*_ + *g*_*s*_(1) < *B* **then**:

7 *x*_*i*_ ← 1

8 *U*← *U* + *s*_*i*_ + *g*_*s*_ (1)

9 **else if** *U* + *s*_*i*_ < *B* **then**

10 *x*_*i*_ ← 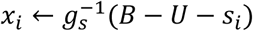 (*B* – *U* – *s*)

11 *U* ← 0

12 **break**

We consider multiple ranking strategies to determine 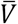 Each control strategy tested is presented in Table 1, as well as the Baseline (B), in which no control is implemented. The first control strategy, Largest Population (LP) simply targets control at airports in the largest cities. The next three strategies exploit known properties of the air traffic network, specifically its hub and spoke structure. The Most Travelled (MT) strategy seeks to target control at airports that are highly trafficked based on total incoming and outgoing volumes, while the Most Connected (MC) and Effective Path (EP) strategies seek to target control at airports that are highly connected to the outbreak source based on travel volumes. The last two strategies utilize learned outcomes from a stochastic epidemic simulation model to inform outbreak control. The First Case (1C) strategy aims to control the set of airports most likely to see the first infected passenger (for a given outbreak scenario), while the First Order Uniform (1OU) targets control at the airports where it is likely to have the largest relative marginal impact. The four strategies, MC, EP, 1C and 1OU utilize knowledge of initial outbreak conditions, while MT and LP airport sets are selected independent of the outbreak state.

**Table 1.** List of control strategies and their description.

| Strategy (Abbreviation) | Method of ranking airports |
| --- | --- |
| Baseline (B) | No control |
| Largest Population (LP) | Airports are ranked in descending order based on the population of the city they serve. When one city is serviced by multiple airports, those airports are further ranked in descending order based on travel volume. |
| Most Travelled (MT) | Airports are ranked in descending order based on travel volume. Travel volume is defined as the sum of daily incoming and outgoing flows. |
| Most Connected (MC) | Airports are ranked in descending order based on total incoming flow from source node(s). |
| Effective Path (EP) | Airports are ranked in ascending order based on their minimum effective path distance from the source(s). For a given outbreak scenario, the effective path distance for each airport is defined as the length of the shortest path from the source, where the length of a path is the sum of the effective distance of each link included in the path. The effective distance of link |
| | $(i, j)$ is defined as $d_{ij} = 1 - \log \left( \frac{\sum_{k \in V} f_{ji}^k(t)}{\sum_{k \in V} \sum_{m \in \Gamma_i^-} f_{mi}^k(t)} \right)^{15}$. When there is more than one source, the minimum effective path distance over all sources is assigned to each airport. |
| First Case (1C) | For a given outbreak scenario the baseline case is simulated a fixed number of times (*i.e.*, 200), and for each run we record the time at which the first infected individual reached each airport, *i.e.*, the initial infection time. Airports are then ranked in ascending order based on their most frequently observed initial infection time. Ties are broken based on the MT criterion. |
| First Order Uniform (1OU) | For a given outbreak scenario the effect of fully controlling each airport independently, *i.e.* $x_i = 1$ for a selected node $i \in V$ and $x_i = 0$ otherwise, is simulated, and the reduction in cumulative number of infected individuals at the observation time $t_{obs}$ is compared to the baseline case (no control). Airports are then ranked in descending order based on the relative reduction in cases, *i.e.*, first-order effects. |

## Data

The metapopulation network is constructed using global passenger air travel data from 2015 provided by the International Air Transport Association (IATA)^54^. The data provided from IATA includes monthly passenger travel volumes for all travel routes connecting airport pairs (including stopovers), representing nearly 83% of global traffic volumes. The final network used in this study contains the top 99% of the travelled routes provided, resulting in a network with approximately 500,000 routes, 2,908 cities, and 3,267 airports. The city populations served by each airport are based on the population densities provided by Oak Ridge National Laboratory’s LandScan^55^. The population size for each city was based on a 50km radius centered on each airport, and computed using open source Geographic Information Systems software QGIS (https://qgis.org/). In some cases, multiple cities are serviced by more than one airport, for which the all assigned airport flows are mapped to the same population.

## Results

We demonstrate the performance of the proposed control strategies to mitigate global outbreak spread, and present results from a cost-benefit analysis, which characterizes the marginal gains in outbreak reduction with respect to increases in available resources, *i.e.,* budget. Further, all strategies are applied to a case study representative of the 2009 H1N1 Pandemic Influenza to illustrate the hypothetical impact of each in a similar outbreak setting. Extensive sensitivity analysis was conducted to assess how the different control strategies respond to differing disease characteristics, model assumptions and policy decisions. Specifically, we explored how changes in the contact rate, planning horizon, control start time, control effectiveness, and source screening impact the performance of each strategy. Results for all sensitivity analysis are provided in the supplementary material (Section A).

In this study, only U.S. cities are considered for control. The two metrics used to compare the performance of each strategy are i) the total cumulative number of infected individuals in the U.S. at observation time, *t*_*obs*_, and ii) the number of infected cities in the U.S. at observation time, *t*_*obs*_ where an infected city has at least one infected individual.

### Base Case Analysis

For the base case analysis, all of the proposed control strategies are implemented and compared for three independent hypothetical outbreak scenarios that vary by the outbreak source location. Three source cities are selected 1) Orlando, Florida, 2) Portland, Oregon and 3) Honolulu, Hawaii, and each is initialized with 100 infected individuals at *t* = 0. These cities were chosen because they represent a range of geographic and travel profiles. In the remainder of this work these three cities are denoted by their assigned airport IATA codes, MCO, PDX and HNL, respectively.

To model the cost of control at airports, we consider linear cost functions. Setup costs *s*_*i*_ represent screening equipment costs based on the total incoming flow at each airport *i* ∈ *V*. Let *M* be the cost of a passenger screening machine and *C* its capacity, we set 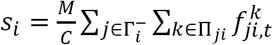. For the variable cost, we assume that the cost of screening a passenger is represented by *P* and set 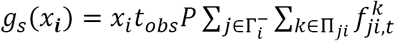 to model the impact of deploying control resources over a varying time period and at a varying level of control. For the base case analysis, we set the available budget *B* to $500 million. The parameter values are *M* = $500,000, *C* = 10,000 and *P* = $10, and based on the existing literature ^56^. Under this configuration, the available budget is enough to fully control the 13 most travelled airports in the U.S. for *t*_*obs*_ =50.

For all scenarios the simulation is set to begin on June 1 and *t*_*obs*_ is set to 50 days. This timeframe aims to model the emerging stage of the outbreak. For the base case analysis, the hypothetical virus has values of α = 0, *β* = 0.25, *γ* = 0.143 and *λ* = 1. The chosen baseline parameters correspond to a disease with a reproductive ratio *R*_0_ = 1.75. All evaluations of the stochastic metapopulation epidemic model are based on 1,000 simulations.

Figure 1 provides the cumulative number of cases in the U.S. at *t*_*obs*_ for scenarios MCO, PDX and HNL. The boxplots capture the results for all 1,000 simulations conducted, illustrating the robustness of the results and rankings. In each plot the six proposed strategies to allocate screening resources are compared against the baseline (corresponding to no control) for the respective scenario. Similar trends are evident for all scenarios, with the more simplistic strategies of controlling at the airports in the largest cities or most travelled airports performing poorest, and EP and MC performing best.

**Figure 1.**
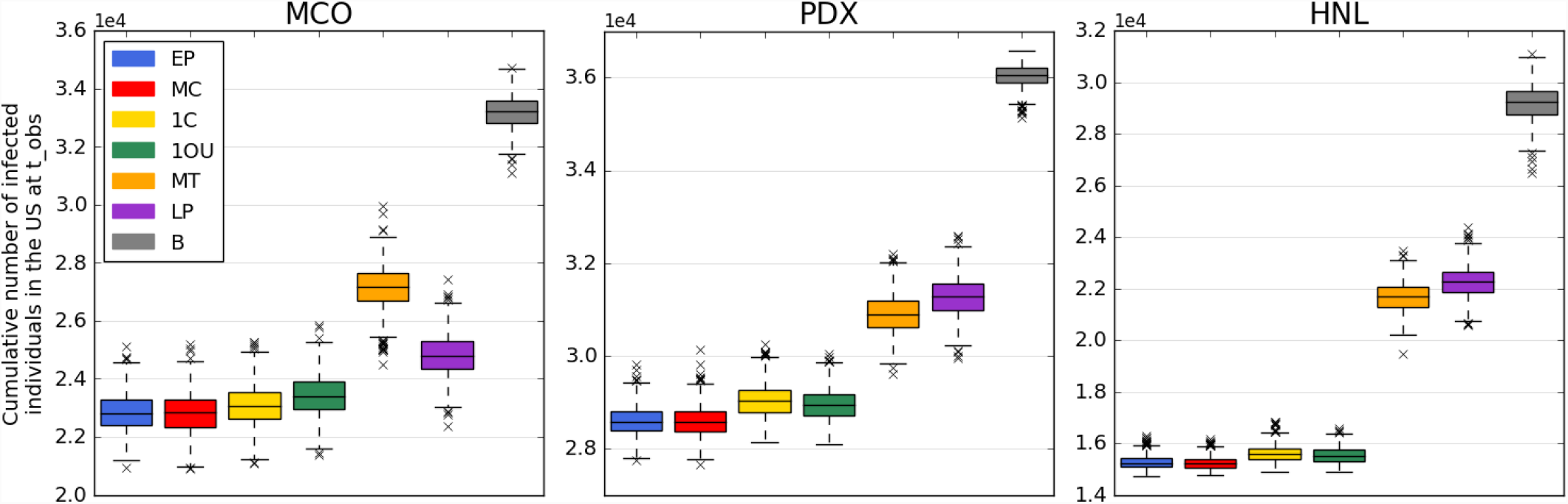
Control strategy performance for the base case scenarios MCO, PDX and HNL in terms of number of infected individuals. The figure reports the cumulative number of infected individuals in the U.S. at the observation time ***t***_***obs***_ = ***50*** days for each control strategy. Each boxplot represents the distribution of the criterion measured over 1,000 simulations of the stochastic metapopulation epidemic model under the corresponding control strategy.

For the MCO scenario, the cumulative number of infected is on average 33,200 for the baseline configuration (no control). Using control strategy LP leads to a reduction of 25.2% in the number of cases, compared with a reduction of 31.2% using EP or MC. For the PDX scenario, the amount of infected individuals is on average 36,000 in the baseline case and using EP or MC results in a reduction of 20.6%. For the HNL scenario, we report a reduction of 47.7% from the baseline to the best control strategy as well as substantially less variability across outbreak simulations.

Figure 2 provides the number of infected cities in the U.S. at *t*_*obs*_ for the three scenarios. This metric is critical to assess the success of the control strategy with regards to preventing spread into new cities. A similar trend is observed across all scenarios, with strategies EP, MC, 1C and 1OU proving superior to MT and LP. For the MCO scenario, the number of cities affected is on average 137 for the baseline case. Using control strategy MT leads to a reduction of 24% in the number of cities affected, compared with a reduction of 31% for EP/MC. For the PDX scenario, the amount of infected is on average 107 in the baseline case and 40 for the EP/MC cases resulting in a reduction of 63%. For the HNL scenario, we observe a more substantial reduction of 90% in the number of cities affected from the baseline to the best control strategy.

**Figure 2.**
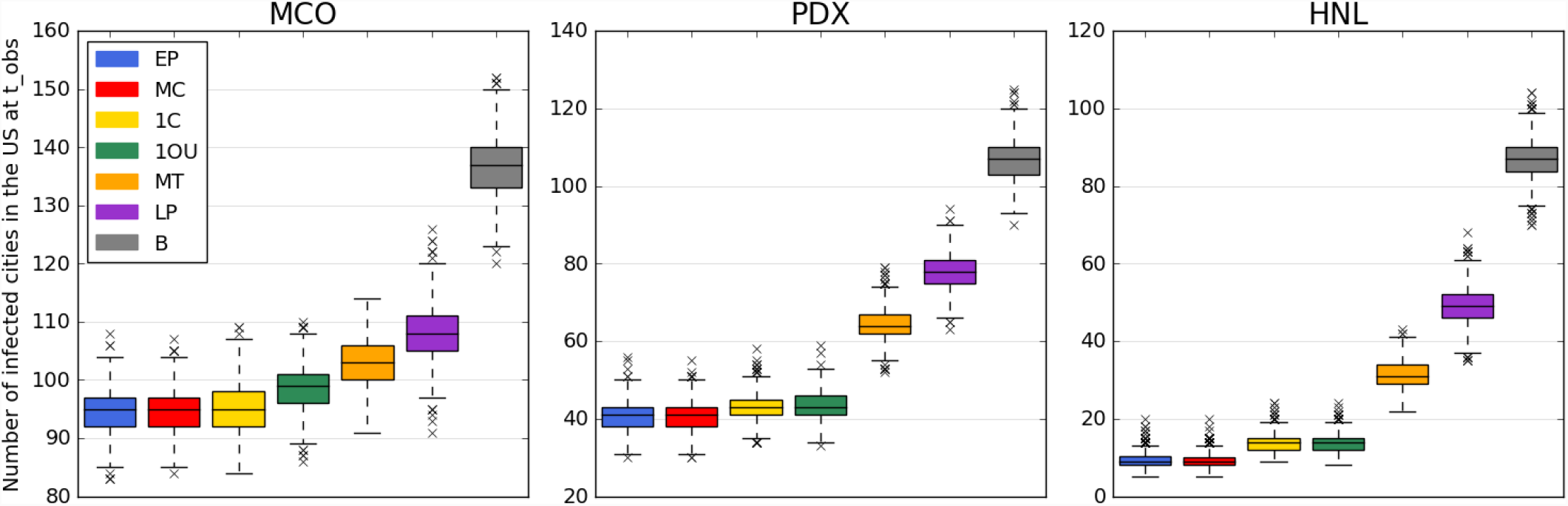
Control strategy performance for the base case scenarios MCO, PDX and HNL in terms of number of infected cities. The figure reports the number of infected cities in the U.S. at the observation time ***t***_***obs***_ = ***50*** days for each control strategy, where a city is categorized as infected if it contains at least one infected individual. Each boxplot represents the distribution of the criterion measured over 1,000 simulations of the stochastic metapopulation epidemic model under the corresponding control strategy.

To help visualize the decisions and impact of the proposed border control strategies, we contrast the outcome of all control strategies with the baseline behavior in Figure 3. The maps display the set of airports selected for control (blue crosses), and the expected size of local outbreaks in each affected city at the observation date (red circles) for each of the five strategies as well as the no-control scenario. The panels correspond to the following strategies: A) No-control, B) MC/EP, C) 1C, D) 1OU, E) MT and F) LP, and the results are illustrated for the PDX scenario. All maps were generated using open source shape files from Natural Earth (http://www.naturalearthdata.com/). Note: EP and MC have the same airport control set in the PDX scenario. The red circles are sized proportional to the outbreak size at each location. The small grey circles represent the set of airports that can be selected for control, further highlighting the scale and the complexity of the problem. Details on the sets of airports selected for each base case scenario and control strategy are provided in the supplementary material (Section C).

**Figure 3.**
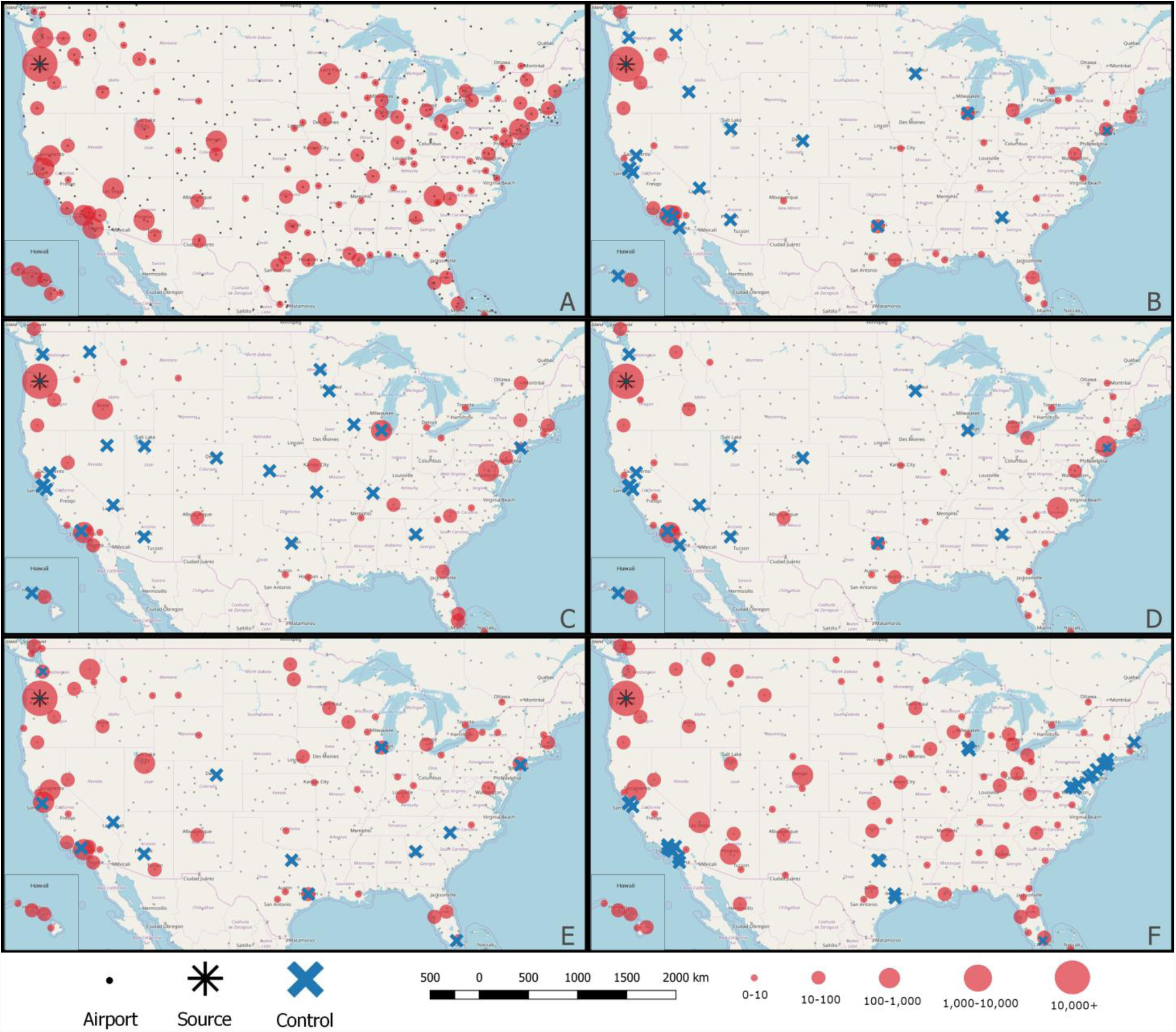
Visualization of outbreak control strategies and impact in the U.S. The results are illustrated for the PDX scenario, with an asterisk marking the source of infection. The maps therein display the set of airports selected for control (blue crosses) for each strategy, and the expected size of outbreak in each city at the observation date *t*_*obs*_ = 50 days (red circles) measured over 1,000 simulations of the stochastic metapopulation epidemic model. The panels correspond to the following strategies: A) No-control, B) MC/EP, C) 1C, D) 1OU, E) MT and F) LP. The red circles are sized proportional to the outbreak size. The grey circles represent the complete set of airports that can be feasibly selected for control. The maps were generated using open source shape files from Natural Earth (http://www.naturalearthdata.com/).

The results indicate both the large spatial variability in the airport sets selected for control across strategies, and the respective impact on outbreak location and outbreak size for each strategy. For example, EP and MC are shown to reduce both the outbreak size and number of affected cities the most at the observation date. The poorest performing strategy is LP, which controls more airports than other strategies, however many of which are far-removed from the outbreak source in terms of traffic volumes, and are therefore less likely to play a major role in furthering spread early in the outbreak. The second worst performing strategy, MT, often spends large portions of its budget on controlling heavily travelled airports, thus resulting in fewer airports controlled, which may also not be critical for outbreak mitigation. It is important to note that both LP and MT airport control sets are fixed for a given budget, and independent of the outbreak scenario, unlike the proposed network- and simulation-driven control strategies.

### Cost-Benefit Analysis

To explore the impact of bresource availability on epidemic spread, we conduct a cost-benefit analysis by varying the budget available for control. Specifically, we explore a range of budgets from $0.25 bil to $1.25 bil, in $0.25 bil increments. Note that given the cost functions used in this study, a budget of $1.35 bil is enough to fully control all airports within the U.S. for a 50-day period. The results are summarized in Figure 4, which illustrates the impact of budget on the effectiveness of each strategy for all three source scenarios. At $0 bil and $1.25 bil the strategies perform nearly the same, which represent the cases of no control and close to full control at all airports, respectively. For budgets in between these values, the impact of all strategies decreases with budget as expected, with control strategy EP consistently outperforming other strategies. The weaker strategies, MT and LP, decrease in a more linear fashion compared to the other four, MC, EP, 1C and 1OU, which show evidence of decreasing marginal returns, *e.g.*, there is negligible improvements after the budget reaches $0.75 bil. These results have valuable implications for policy makers, and indicate the potentially cost-effective nature of resource allocation if assigned strategically.

**Figure 4.**
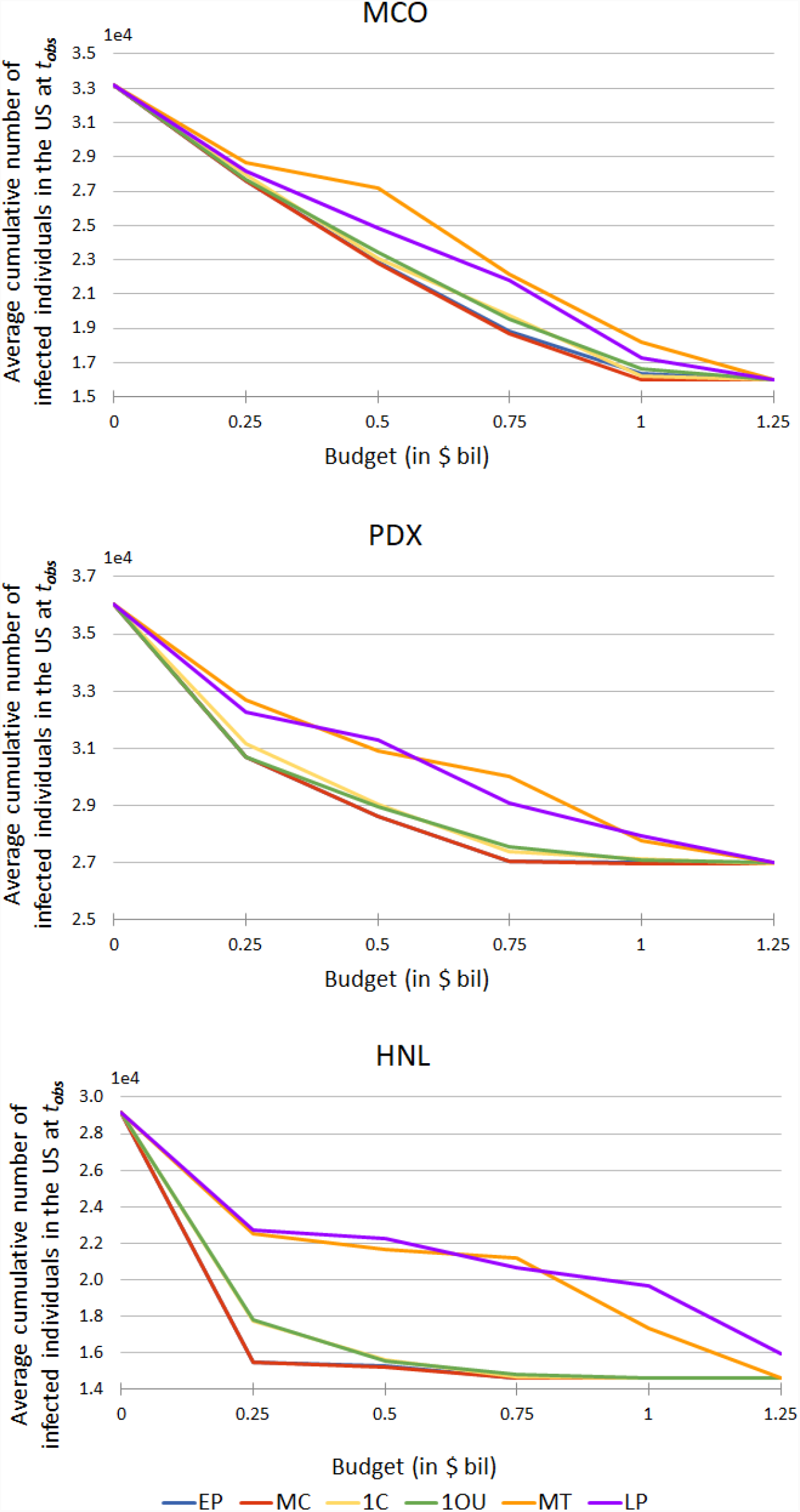
Cost-benefit analysis for the base case scenarios MCO, PDX and HNL in terms of number of infected individuals. The figure depicts the average cumulative number of infected individuals in the U.S. at the observation time ***t***_***obs***_ = ***50*** days based on a varying border control budget *B* expressed in billions of dollars. The data points converge at *B* = **0** which corresponds to the baseline (no control) case.

### Case Study: 2009 H1N1 Influenza Pandemic

To illustrate the ability of the simulation model to replicate a realistic pandemic, we consider the 2009 H1N1 influenza pandemic as a case study, and quantify the performance of the proposed strategies under similar outbreak conditions. The simulation model was calibrated to the 2009 H1N1 outbreak data at both the U.S. and global scales. The calibration methodology and results are included in the supplementary material (Section B). The final calibrated disease parameters are *α* = 1, *β* = 0.475, *γ* = 0.25 and *λ* = 1. For all simulations the starting date of the outbreak is set as the 5^th^ of February 2009 with 1 infected individual placed into the city of Veracruz, Mexico^1^. Each control strategy is deployed four weeks (28 days) after the first case appeared in Mexico, at which time there were about 100 local cases in Mexico (based on the simulation). The cost function, budget and screening costs are the same as those used in the base case analysis. The results are based on 1,000 simulations of the stochastic model, and the observation time is set to *t*_*obs*_ = 100 days after the first case.

The case study illustrates the hypothetical impact of implementing the proposed strategies for an outbreak similar in characteristics to the 2009 H1N1 influenza pandemic. The performance of each strategy based on the two metrics used for evaluation, *i.e.*, the cumulative number of cases and number of infected cites in the U.S. at the time of observation, under each of the proposed strategies are illustrated in Figure 5. The results highlight the EP strategy to once again dominate, and LP and MT to perform the poorest. Specifically, the EP strategy can reduce the final outbreak size by 37.1% relative to the baseline, with the number of infected cities dropping from 115 to 86, representing a 25.2% decrease.

**Figure 5.**
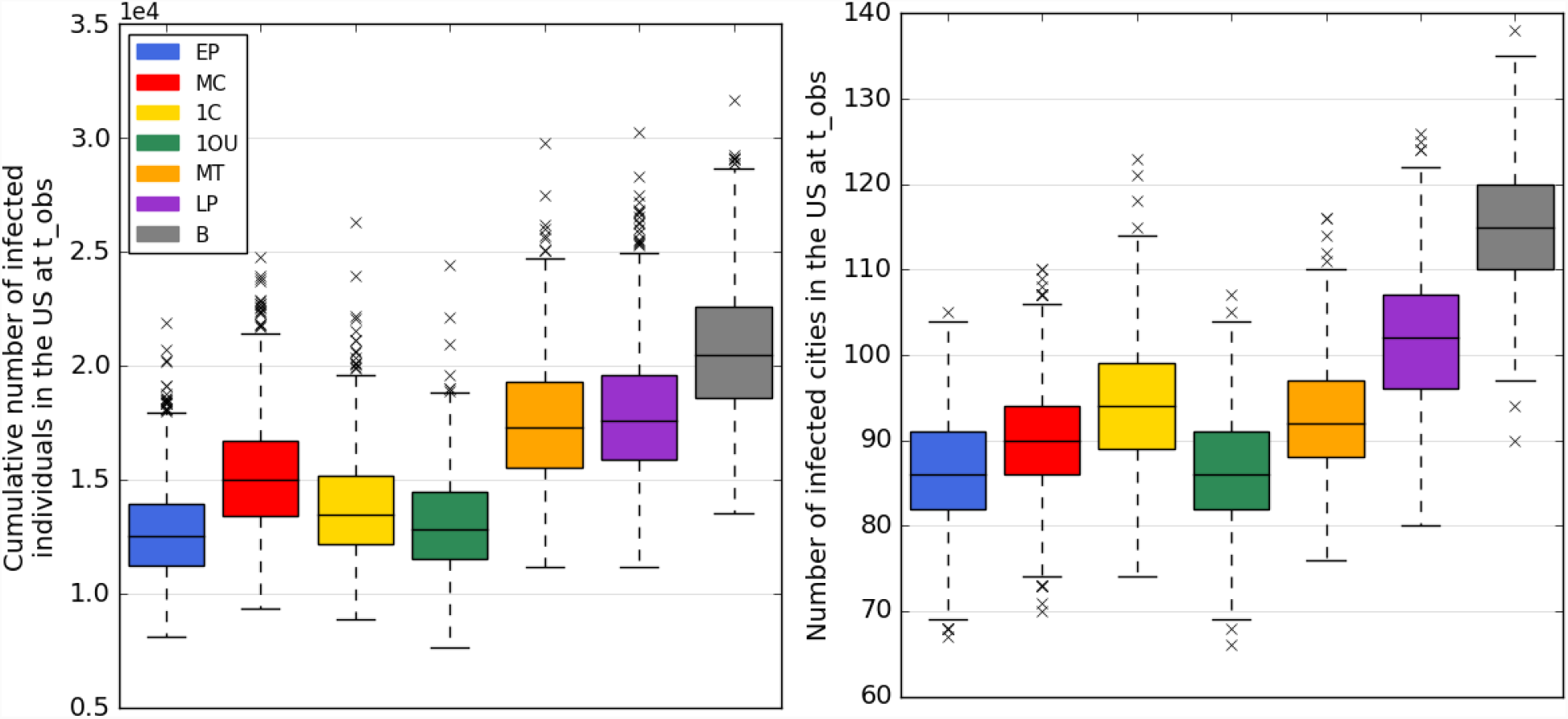
Control Strategy performance for the 2009 H1N1 influenza pandemic case study in terms of number of infected individuals (left) and number of infected cities (right) in the U.S. The figure depicts the performance of each control strategies for an observation time ***t***_***obs***_ = **100** days, and assuming border control is deployed at **28** days after the first infected individual was reported in Mexico. Each boxplot represents the distribution of the criterion measured over 1,000 simulations of the stochastic metapopulation epidemic model under the corresponding control strategy.

## Discussion

This work addresses the challenge of pandemic mitigation planning through border control, specifically using entry screening at airports to minimize the potential harm posed by an outbreak. We present an ensemble of control strategies that are evaluated based on their ability to reduce the cumulative number of cases and the number of cities infected at a target observation time. The decision-support framework provided can be implemented in real-time at the early stages of a confirmed or suspected outbreak. An in-depth analysis is presented for multiple hypothetical outbreak scenarios, and a case study is conducted to illustrate the performance of the control strategies for an outbreak similar to the 2009 H1N1 influenza pandemic. Further, extensive sensitivity analysis illustrates the robustness of the control strategies to various modelling parameters and assumptions.

The best performing control strategies are the network-driven strategies, *e.g.*, EP and MC, which are shown to be robust across a range of outbreak scenarios and model assumptions. In contrast, the more simplistic strategies, *e.g.*, controlling the most travelled airports (MT) or the airports in the largest cities (LP), perform the poorest. The superiority of the network-driven strategies highlights the significance of the heterogeneity of the world air traffic network in outbreak spreading^5,15^, which can be exploited by policy makers for the purposes out pandemic planning and mitigation. The two control strategies informed by simulation, 1OU and 1C, rank in the middle in terms of performance in most scenarios considered in this work, however the relative performance of the strategies is sensitive to the outbreak initial conditions, as highlighted by the case study. While the results from the analysis presented indicate the EP strategy to be reliably superior, the larger contribution of this work is the modeling framework, which can be implemented in real-time for any outbreak scenario given reported case counts and locations. Additionally, the integrated framework is flexible, and can be easily extended to incorporate and compare the performance of additional strategies if desired.

The cost-benefit analysis highlights two critical issues. Firstly, a given budget can be used more effectively if the control decisions are made strategically. *e.g.*, EP/MC can achieve the same effectiveness as MT/LP for nearly half the budget. Second, there is evidence of decreasing marginal returns for the superior strategies, indicating minimal benefits may be gained by spending more on border control beyond a certain threshold. This cost-effectiveness threshold is critical for policy makers, who can choose to redirect available control resources towards alternative control strategies for outbreak mitigation which may be more effective.

The case study illustrates the expected performance of each control strategy for an outbreak similar in behavior to the 2009 H1N1 influenza pandemic. This analysis critically introduces an asymptomatic exposed state, which poses an additional challenge for control, because even with perfect screening not all infected travelers can be identified. Even so, results from the case study reveal the reliably superior performance of EP as an outbreak control strategy, and once again, the value gained by strategically allocating control resources.

The sensitivity analysis on the infection contact rate (see Section A.1) illustrates that the ranking and effectiveness of the control strategies are robust with regards to the rate of spread. Furthermore, the control strategies are observed to be more effective (relative to the no control scenario) and more robust for faster spreading viruses. The more reliable performance for higher contact rates can be attributed to the highly stochastic nature of outbreaks in their early stages, which is heavily dependent on where the first few infected cases spread to, whereas a faster spreading virus will result in a larger number of infected people travelling, reducing the variability across outbreak scenarios.

Sensitivity to observation time and control start date was conducted to illustrate the impact of these two implementation options available to policy makers. The model appears to be highly sensitive to observation time (see Section A.2), which is due to the exponential nature of outbreak growth; there is a small number of cases and affected cities in the early stages of the outbreaks (*e.g.* 25 days) compared to considerably higher infection rates at later time epochs (*e.g.* 100 days). This sensitivity analysis highlights the critical (short) timeline during which border control has the potential to play a substantial role, after which local control will be most impactful. The impact of delaying border control was also evaluated (Section A.3), and the results again highlight the robustness of the strategy rankings. The results also demonstrate the importance of implementing control in a timely fashion, with the best performing strategies revealed to be the most sensitive to delayed start times.

To address the assumption of perfect control, we evaluated the effectiveness of the proposed strategies under imperfect control conditions (Section A.4), limiting the maximum control rate to 80% and 90%, respectively. While the impact of control decreases with the control level, the relative performance and ranking across strategies remains constant, suggesting the best performing strategies remain effective and reliable under imperfect control. Similar results are observed in the case study, which utilizes an SEIR model, *i.e.*, asymptotic infected travelers are able to evade control and introduce infection into new cities.

The final sensitivity analysis evaluated the impact of outgoing passenger screening at the source of infection. This work assumes that outgoing screening at the source is an obvious decision; therefore the model addresses the more challenging problem of selecting which locations other than the source(s) should be prioritized for incoming passenger screening. The sensitivity analysis results reveal that when outgoing screening is implemented at the source (Section A.5) the proposed control strategies behave predictably, *i.e.,* the outbreak spreads faster but the strategy rankings remain consistent. Critically, the best performing strategies perform well even at low levels of outgoing screening, *i.e.*, ineffective screening, while the poorest performing strategies are more sensitive to the effectiveness of outgoing source screening.

Lastly, there are modeling assumptions and limitations of this study. First, the model only accounts for passenger air travel, and excludes mobility within and between cities via other modes of transport. Second, local disease spread (within a city) is modeled deterministically, and a uniform contact rate is used across all populations. Third, the model is currently limited to global control decisions through passenger screening, and does not evaluate local control mechanisms (prophylaxtics, vaccines, school closures, etc), with the exception of source exit screening. Fourth, airport (rather than route) screening rates are the control variable, and therefore assume passenger screening to be uniformly applied across all incoming routes at a given airport. Planned extensions of this work will address these limitations through i) integrating alternative modes of transport into the model, ii) adding additional decision variables to optimize local control decisions, iii) the development of a link-based modelling formulation to allow specific travel routes to be identified for screening (as opposed to airports), and iv) introducing a dynamic resource allocation formulation which relaxes the assumption of constant control across the entire planning period. These extensions would provide more degrees of freedom to improve the impact of control resources and further help in minimizing the risk posed by global outbreaks. The set of limitations and extensions listed currently lie outside the scope of this study, and provide the basis for future research.

## Data Availability

The air traffic data used in this study is available for purchase from IATA Passenger Intelligence Services (PaxIS), https://www.iata.org/services/statistics/intelligence/paxis. The population data is publically available from ORNL’s LandScan, https://landscan.ornl.gov/. H1N1 case data used for model calibration is publically available from CDC and WHO, and referenced in the supplementary materials.

## Author Contributions

The authors contributed to (A) conceive and design the experiments, (B) perform the experiments, (C) write the paper, (D) develop the model, (E) perform the data driven simulations and (F) analyse the data. Aleksa Zlojutro B, C, D, E, F; David Rey A, C, D, F; Lauren Gardner A, C, D, F. All authors reviewed the results and approved the final version of the manuscript.

## Additional Information

The author(s) declare no competing interests

## Supplementary Material

## A. Sensitivity Analysis

To investigate how the strategies respond to the various parameters and model assumptions, we conduct a range of sensitivity analysis. We explore how changes in the contact rate, the target observation time, *i.e*. *t*_obs_, control start time (delayed control), imperfect compliance, and screening outgoing travelers at the source of infection impact the performance of each control strategy. For this sensitivity analysis the set of source cities and base case conditions remain the same as those described in the main document (see Base Case Analysis)

### A.1 Impact of the Contact Rate, *β*

A significant source of uncertainty in a novel infectious disease is its level of infectiousness, *i.e.*, how fast it will spread. To explore the robustness of our proposed control strategies to the infectivity of a disease, we conduct sensitivity for a range of contact rates. The results for *β* = 0.2, 0.25 and 0.3 are shown for all scenarios MCO, PDX and HNL, in Figures S1a, S1b and S1c, respectively. These three contact rates (CR in the figures) correspond to a disease with a reproductive ratio *R*_0_ = 1.4, 1.75 and 2.1, respectively.

As expected, the number of cases increases with the contact rate for all control strategies. The relative performance of the control strategies compared with the baseline case (corresponding to no control being deployed) increases as the contact rate increases, indicating that the control strategies are more effective for more aggressive outbreaks. Specifically, for the PDX scenario, the EP and MC strategies provide a 16% decrease in cases for *β* = 0.2 compared with a 24.9% for *β* = 0.3, relative to the baseline. The gap between the poorest performing strategy, MT, and the best performing strategy also increases with *β*. Across the range of *β*, EP and MC consistently perform best. These trends are reflected in scenarios MCO and HNL, with the latter exhibiting a significantly lower volatility in terms of performance compared to the other source city scenarios.

**Figure S1a.**
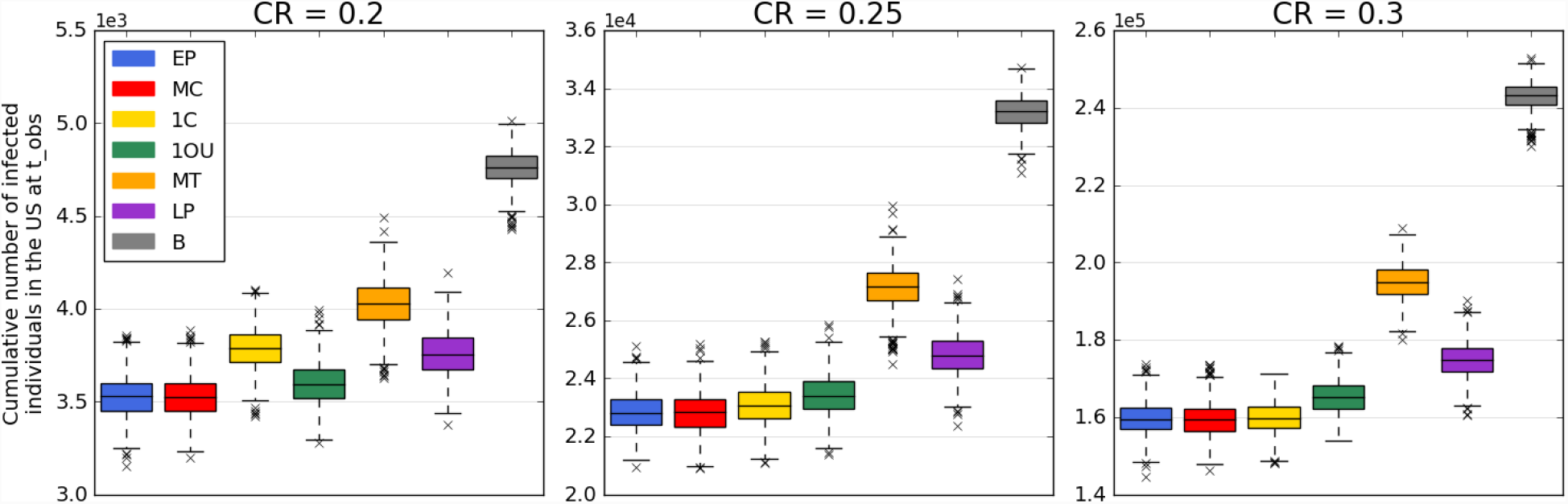
Impact of the contact rate on the control strategies for the MCO scenario. The figure reports the cumulative number of infected individuals in the U.S. at the observation time ***t***_***obs***_ = ***50*** days for each control strategy evaluated based on the contact rate (CR), denoted *β* in the formulation. Each boxplot represents the distribution of the criterion measured over 1,000 simulations of the stochastic metapopulation epidemic model under the corresponding control strategy.

**Figure S1b.**
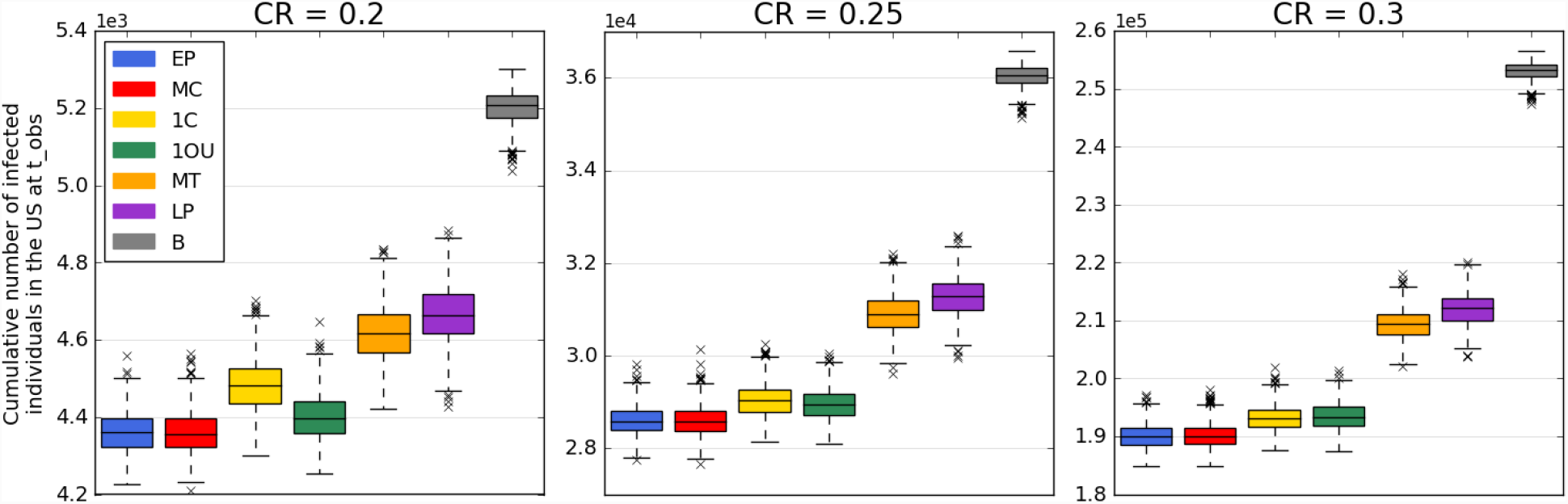
Impact of the contact rate on the control strategies for the PDX scenario. The figure reports the cumulative number of infected individuals in the U.S. at the observation time ***t***_***obs***_ = ***50*** days for each control strategy evaluated based on the contact rate (CR), denoted *β* in the formulation. Each boxplot represents the distribution of the criterion measured over 1,000 simulations of the stochastic metapopulation epidemic model under the corresponding control strategy.

**Figure S1c.**
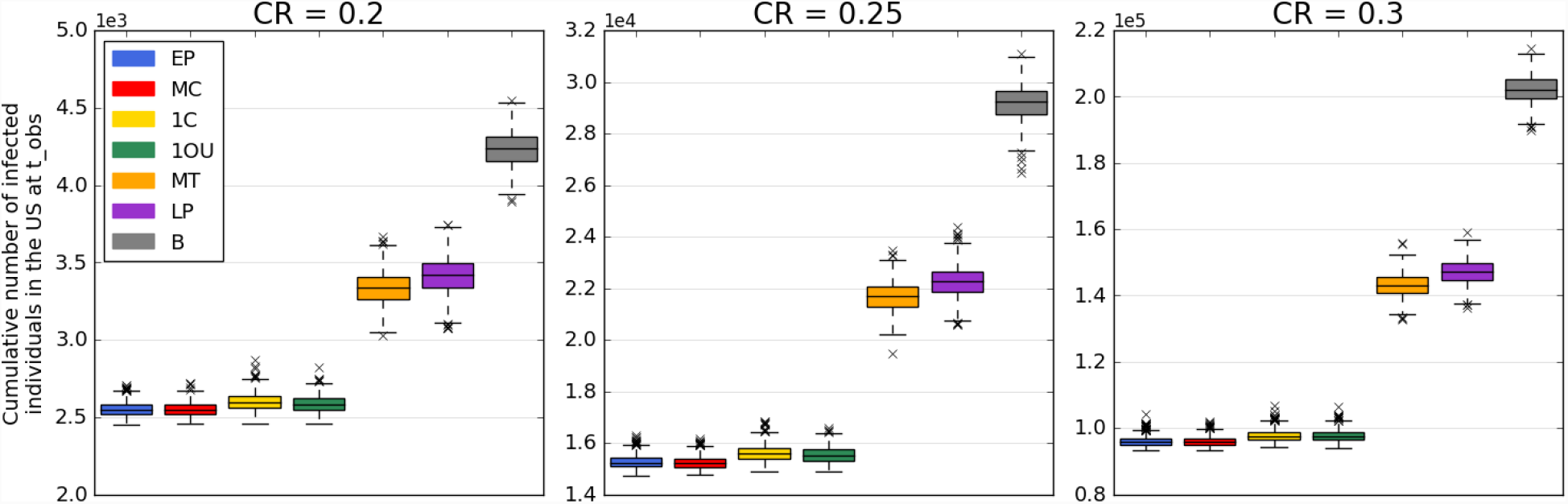
Impact of the contact rate on the control strategies for the HNL scenario. The figure reports the cumulative number of infected individuals in the U.S. at the observation time ***t***_***obs***_ = **50** days for each control strategy evaluated based on the contact rate (CR), denoted *β* in the formulation. Each boxplot represents the distribution of the criterion measured over 1,000 simulations of the stochastic metapopulation epidemic model under the corresponding control strategy.

### A.2 Impact of varying the observation time

The proposed decision-support framework is based on the chosen time of observation, *t*_*obs*_. As previously discussed, the focus of this work is to help guide control decision at the early stages of an outbreak, with the intention of preventing introductions into new cities and regions. Thus, the planning horizon focuses on the first weeks and months after a new disease has been identified. However, the impact of the chosen planning horizon is critical to quantify, as it directly relates to the budget, *e.g*., a longer planning horizon requires more days of screening, and therefore less airports can be controlled for the same budget. A sensitivity analysis is conducted to measure the impact of *t*_*obs*_ when this parameter is varied between 25 and 100 days and its impact on the control strategy performance is assessed. Figures S2a, S2b and S2c provide the results of the sensitivity analysis for scenarios MCO, PDX and HNL, respectively. As *t*_*obs*_ increases, the number of cumulatively infected individual grows quickly, highlighting the non-linear growth of outbreaks. The relative impact of the best control strategies, MC and EP, continues to increase with *t*_*obs*_, and critically, they consistently dominate in terms of performance across planning horizons, indicating the strategies are robust. For the MCO scenario, we find that the control strategies perform increasingly similarly when the observation time increases. This is not the case for the PDX scenario wherein increasing *t*_*obs*_ magnifies the differences of the control strategies in terms of performance. Finally, the HNL scenario is found to be the most robust to the variation of the observation time and this trend is also reflected more generally for this source city scenario.

**Figure S2a.**
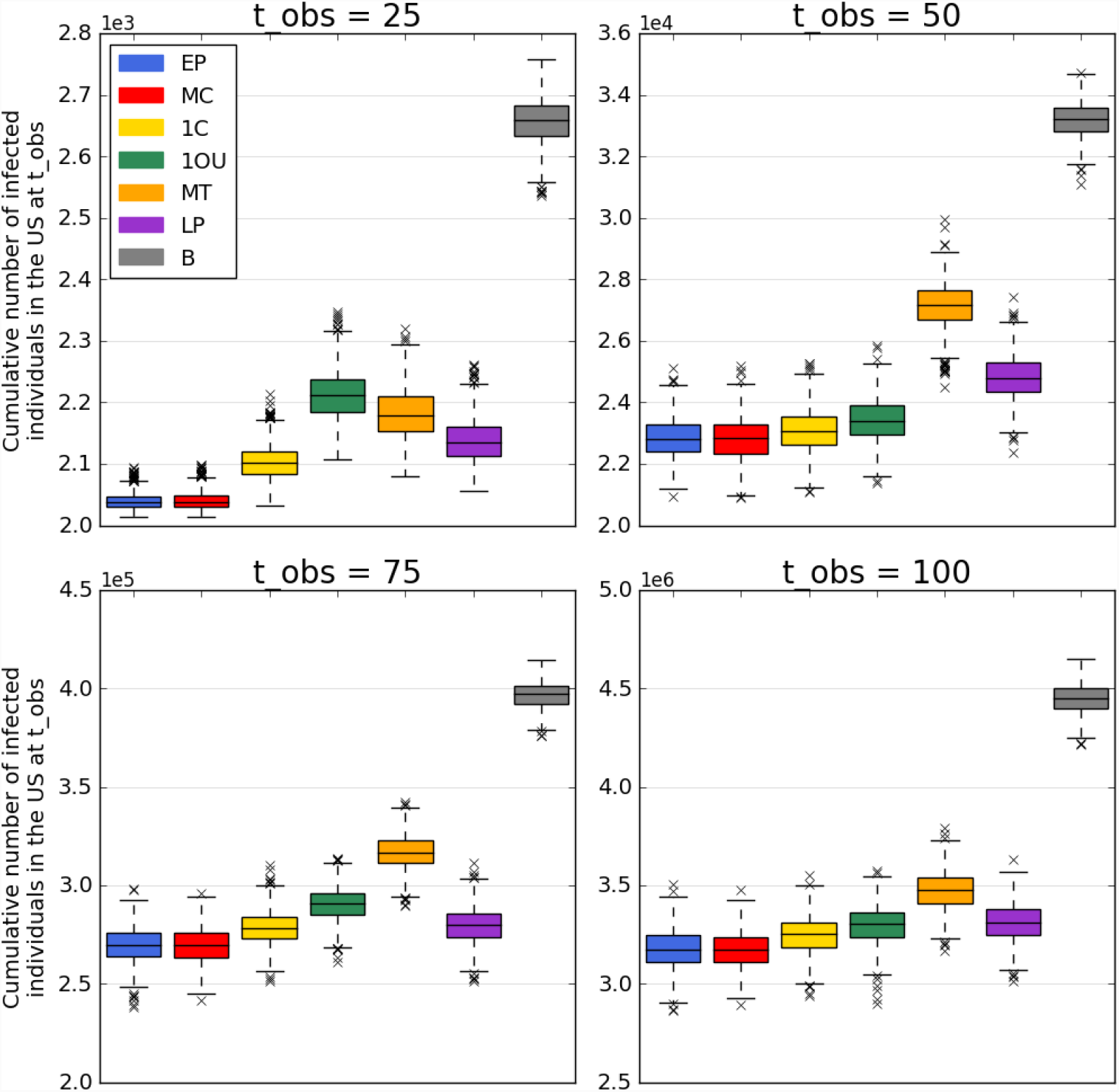
Impact of changing *t*_*obs*_ on the control strategies for the MCO scenario. The figure reports the cumulative number of infected individuals in the U.S. at the observation time ***t***_***obs***_ = ***50*** days for each control strategy evaluated based on the time of observation ***t***_***obs***_. Each boxplot represents the distribution of the criterion measured over 1,000 simulations of the stochastic metapopulation epidemic model under the corresponding control strategy.

**Figure S2b.**
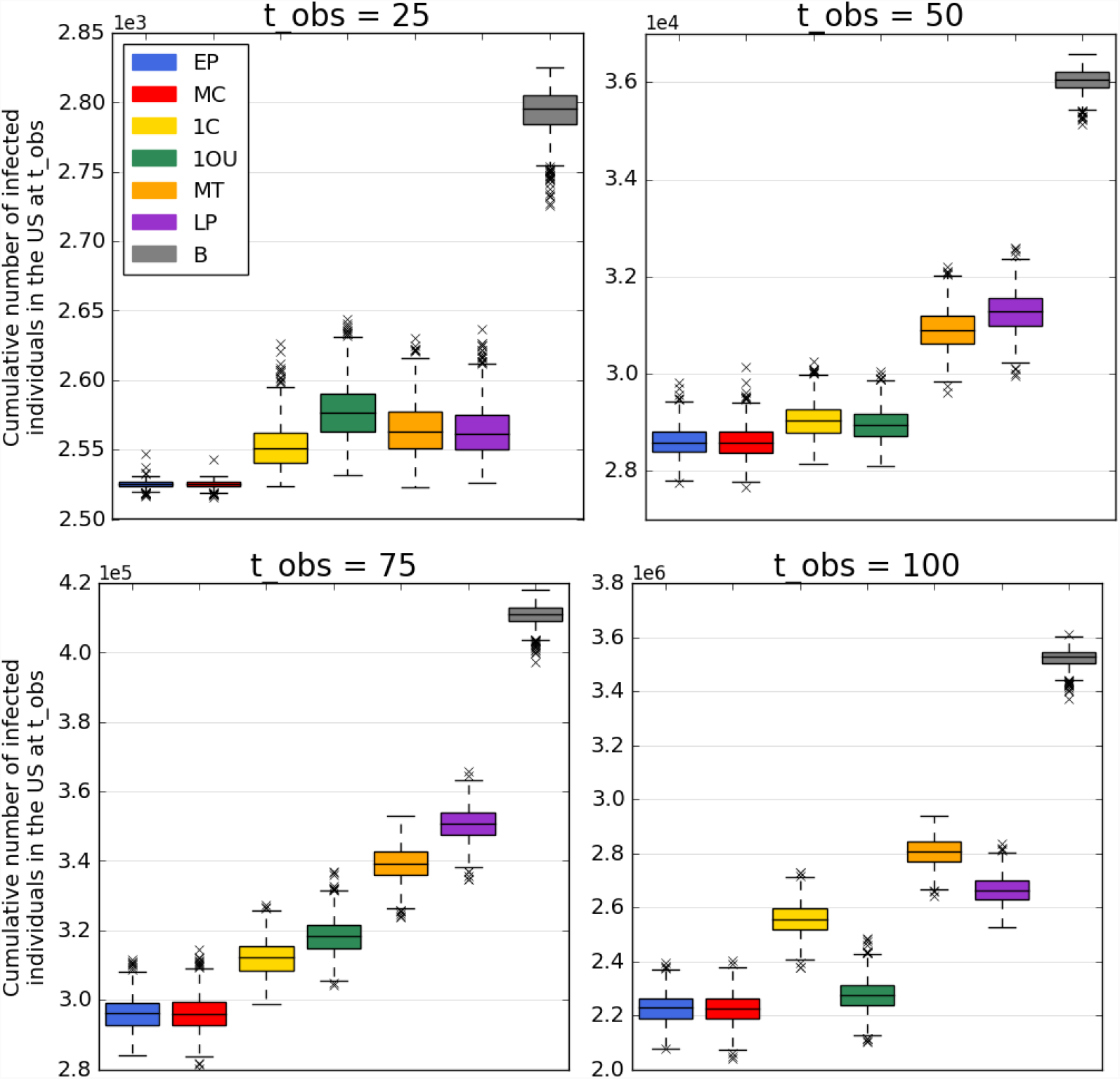
Impact of changing *t*_*obs*_ on the control strategies for the PDX scenario. The figure reports the cumulative number of infected individuals in the U.S. at the observation time *t*_*obs*_ = 50 days for each control strategy evaluated based on the time of observation *t*_*obs*_. Each boxplot represents the distribution of the criterion measured over 1,000 simulations of the stochastic metapopulation epidemic model under the corresponding control strategy.

**Figure S2c.**
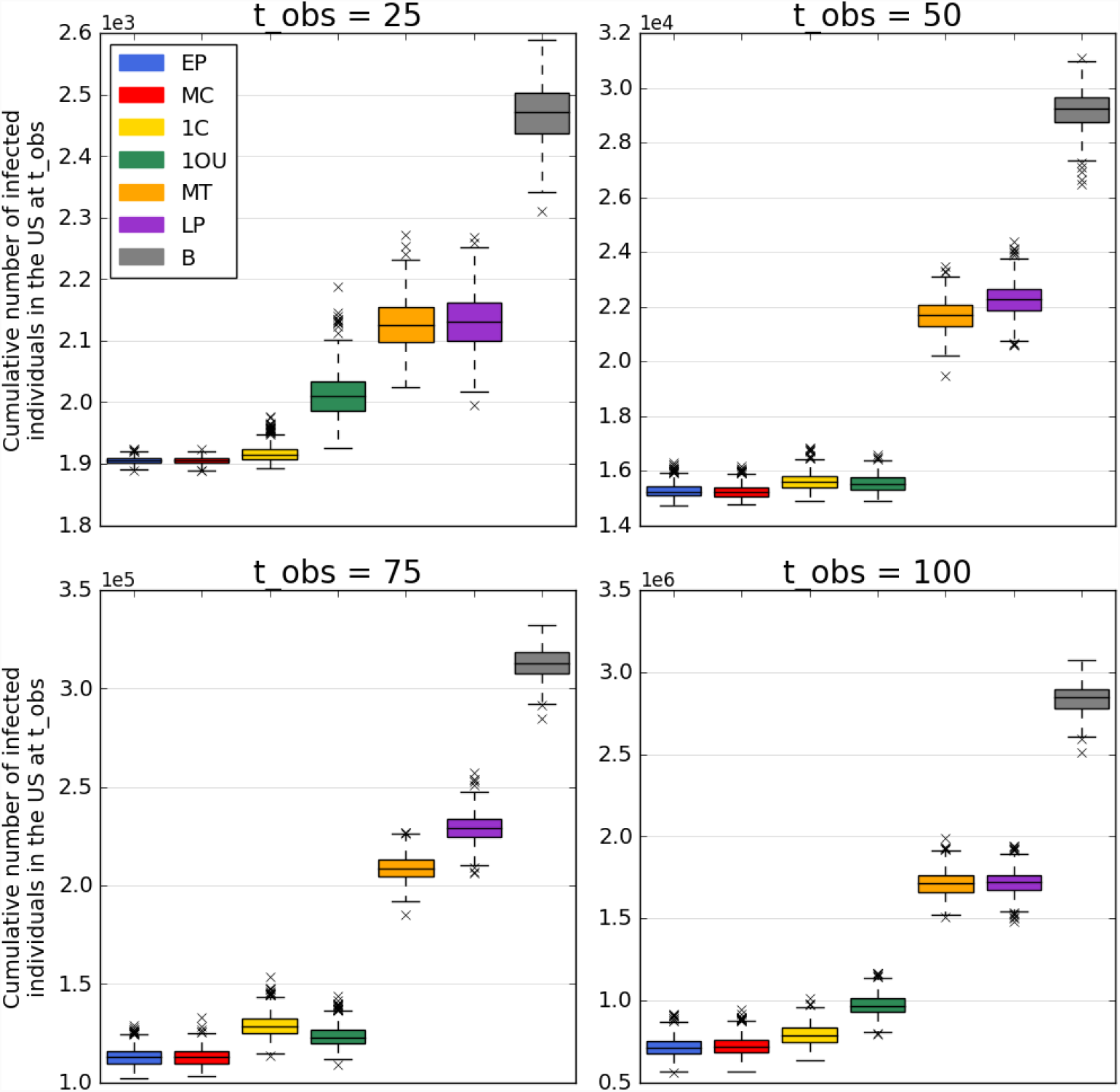
Impact of changing *t*_*obs*_ on the control strategies for the HNL scenario. The figure reports the cumulative number of infected individuals in the U.S. at the observation time ***t***_***obs***_ = ***50*** days for each control strategy evaluated based on the time of observation ***t***_***obs***_ Each boxplot represents the distribution of the criterion measured over 1,000 simulations of the stochastic metapopulation epidemic model under the corresponding control strategy.

### A.3 Impact of Delaying Control

As this work focuses on the early stages of an outbreak, the impact of delaying the deployment of control resources can provide key insights on policy implementation and practice. To this effect, we conduct a sensitivity analysis time at which control, *i.e.*, passenger screening is deployed. Specifically, a delay of 0, 7, 14, 21 and 28 days is evaluated – we note that in the base case analysis, control is assumed to start at *t* = 0 whereas in the case study control is assumed to start at *t* = 28 days. In these scenarios the airports control sets for each strategy remain the same as in the base case, with the only difference being the start date of screening. The results of the scenarios are presented in Figure S3. Note: EP performs closely to MC, and is therefore not clearly visible in the figure.

In all base case scenarios MCO, PDX and HNL, delaying the deployment of passenger screening results in more cases. Further, we observe that all control strategies are similarly impacted and their ranking, in terms of their capacity to reduce the number of infected individuals, remain almost always the same (except in scenario HNL, wherein MC/EP and 1OU crossover). The results also highlight that the smarter control strategies (*i.e.*, EP, MC, 1OU and 1C) are more sensitive to delayed implementation, indicating a critical need to implement control as soon as a risk is identified.

**Figure S3.**
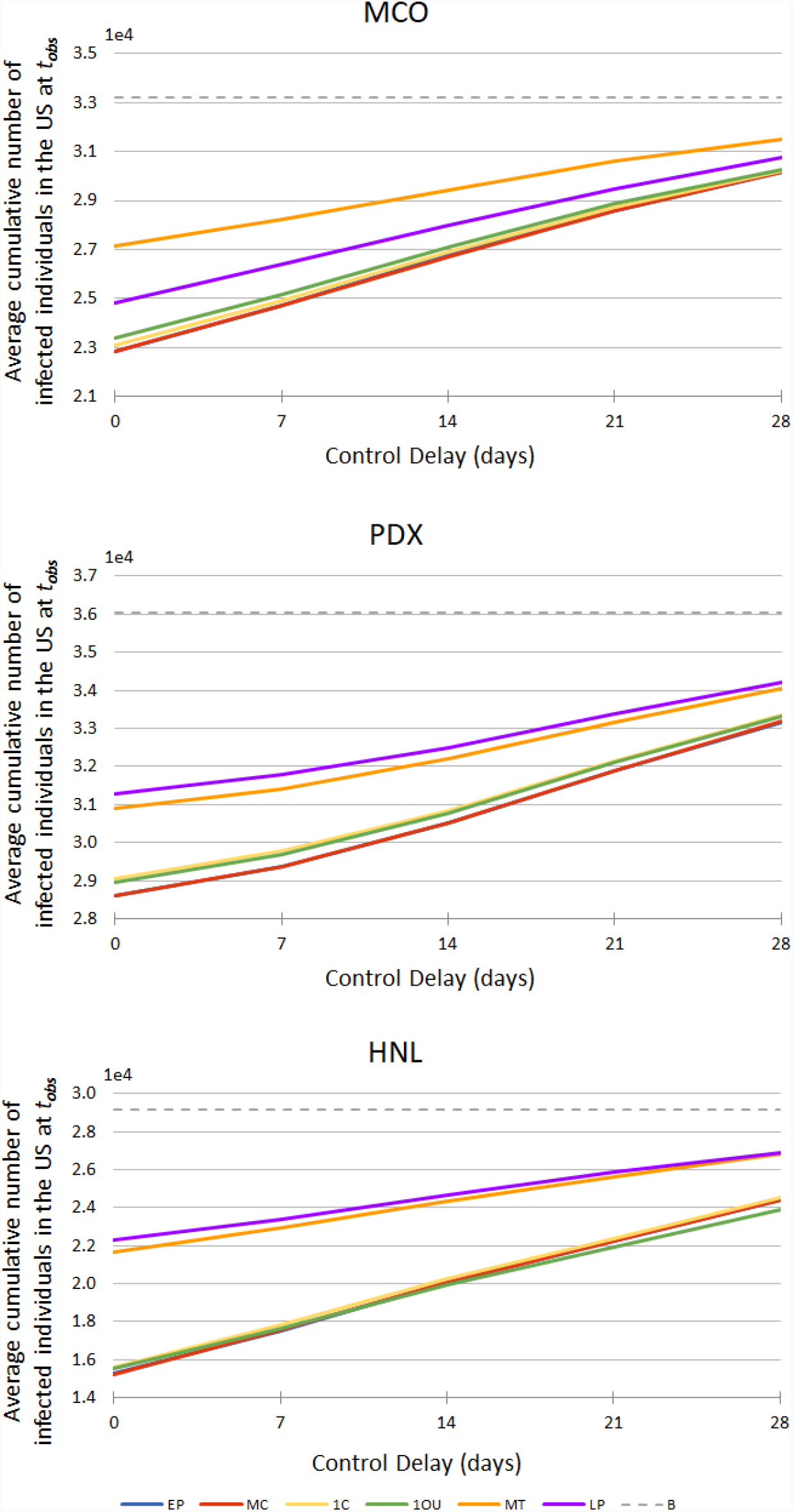
Impact of delayed control the control strategies for base case scenarios MCO, PDX and HNL. The figure reports the average cumulative number of infected individuals in the U.S. for each control strategy evaluated for a varying control start time, *i.e.*, delayed control. Each data point represents the average of the criterion measured over 1,000 simulations of the stochastic metapopulation epidemic model under the corresponding control strategy.

### A.4 Impact of Imperfect Compliance

A sensitivity analysis is also conducted to address the modelling assumption of fully successful passenger screening. Indeed, if airport *i* ∈ *V* is fully controlled, *i.e. x*_*i*_ = 1, our model assumes that all infected individuals are successfully detected in the screening process. In reality, this may not be the case. For example, Auckland International Airport’s screening procedure for the 2009 influenza pandemic found that only 6% of infected passengers were detected^1^. This was partially due to the inability of the airport staff to screen all incoming passengers, as well as the limitations of the thermal scanning technology that was used. It is therefore necessary to evaluate the effectiveness of the proposed strategies under imperfect control conditions. To explore the impact of imperfect control, we set upper bounds lower than 1 on the control rate at airports and measure performance in the base case scenarios. The outcome is reported Figure S4. The performance of each control strategy is presented for the cases where the control rate is limited to 0.9 and 0.8. The results are compared with the perfect control scenario. We assume that the same set of airports is controlled (obtained from the base case outcomes) in all simulations.

The results once again highlight the robustness of the control strategies. The increase in infected cases increases as the perfect control assumption is relaxed, however the relative performance and ranking across strategies remains robust to this relaxation. We also report a more apparent increase in cases between full control and 90% control, compared with the difference in 80% and 90% control effectiveness.

**Figure S4.**
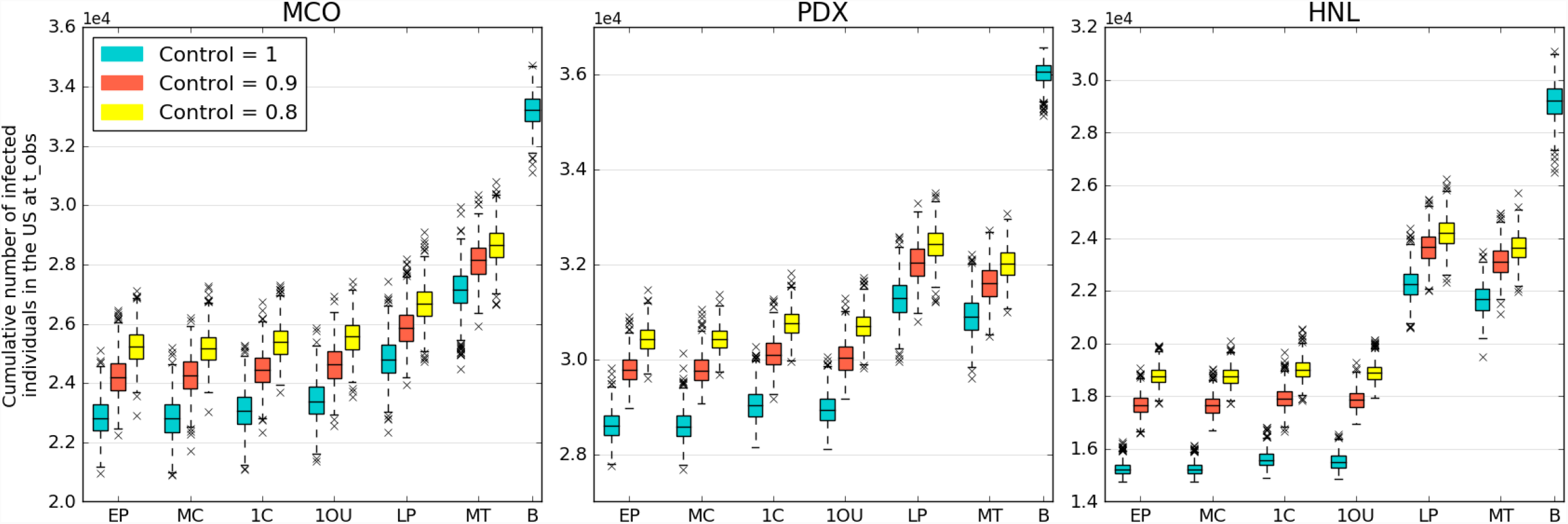
Impact of imperfect compliance on the control strategies for base case scenarios MCO, PDX and HNL. The figure reports the cumulative number of infected individuals in the U.S. for each control strategy evaluated for a varying control start time, *i.e.*, delayed control. Each boxplot represents the distribution of the criterion measured over 1,000 simulations of the stochastic metapopulation epidemic model under the corresponding control strategy.

### A.5 Impact of Screening Outgoing Travelers at the Source of Infection

A final sensitivity analysis addresses the issue of outgoing passenger screening at the source of the outbreak. In this work we assumed that screening outgoing passengers at the source city of the outbreak is a trivial decision. Thus, the proposed control strategies focus on which ‘other’ locations should be prioritized for incoming passenger screening. Furthermore, if outgoing passenger screening is conducted at the outbreak source city, assuming a uniform effectiveness across all intended destinations, the relative performance proposed strategies still applies. This is illustrated in Figure S5, which provides the different control strategy performances and ranking for source screening rates of 0%, 25%, 50%, 75%, 100%. At a 100% outgoing screening rate, no cases will ever leave the source city, thus all strategies perform equally, while the 0% outgoing screening rate corresponds to the base case results. A linear response to source screening is observed for all strategies as the level of source screening is varied. Once again, the strategy rankings remain consistent across scenarios. In all cases, the cost of screening outgoing passengers is excluded from the budget.

**Figure S5.**
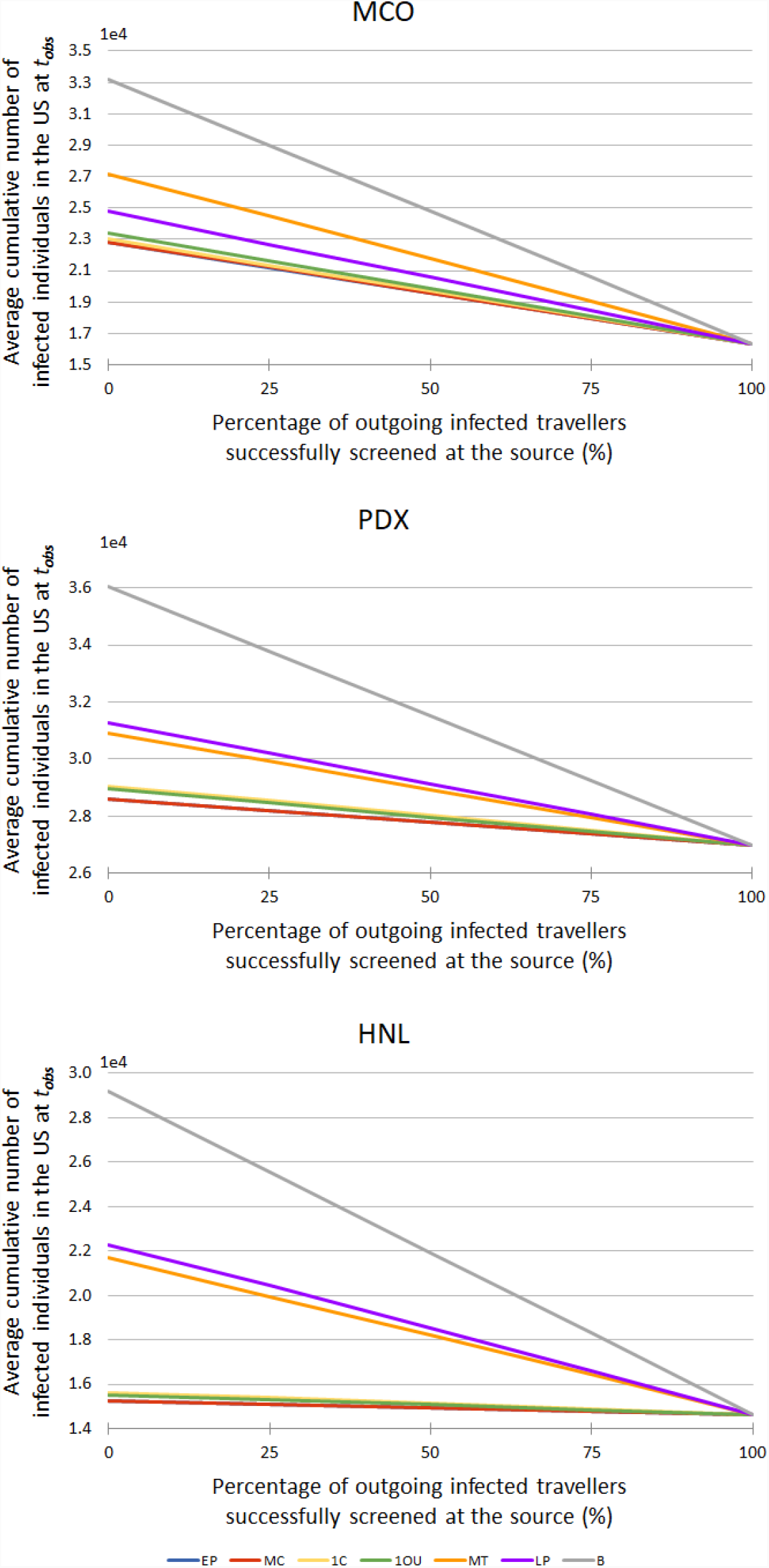
Impact of screening outgoing travelers at the source of infection on the control strategies for base case scenarios MCO, PDX and HNL. The figure reports the cumulative number of infected individuals in the U.S. for each control strategy evaluated for a varying proportion of outgoing infected travelers successfully screened at the source city. Each data point represents the average of the criterion measured over 1,000 simulations of the stochastic metapopulation epidemic model under the corresponding control strategy.

### B. H1N1 Case Study Model Calibration

In this section, we provide additional details on the calibration of the proposed stochastic metapopulation epidemic model. The focus of the calibration process was on matching the simulated and reported arrival dates of the first case in a new location, at both the U.S. and global scales. The parameters of the model were set such that the average difference between the reported date of first case^2,3^ and the simulated date (when averaged over 1,000 simulations) was minimized. For all simulations the starting date of the outbreak is set as the 5^th^ of February 2009 with 1 infected individual placed into the city of Veracruz, Mexico^4^. Based on the analysis, the best fit disease parameters were found to be α = 1, *β* = 0.475, *γ* = 0.25 and *λ* = 1.

Figure S6 compares the observed and simulated date of H1N1 introduction into each of the states in the U.S. for the calibrated model. The model is illustrated to closely capture the trend of first case introductions; critically capturing the gap between the start of the outbreak in Mexico and the time of introduction into California and Texas, after which the outbreak quickly spread to the rest of the country. For the chosen parameters, the average difference between the simulated and observed date of arrival averaged across all 50 states is 7.28 days. The CDC reported dates are later than the dates of initial infection and hence slightly earlier prediction dates by the model are preferable.

**Figure S6.**
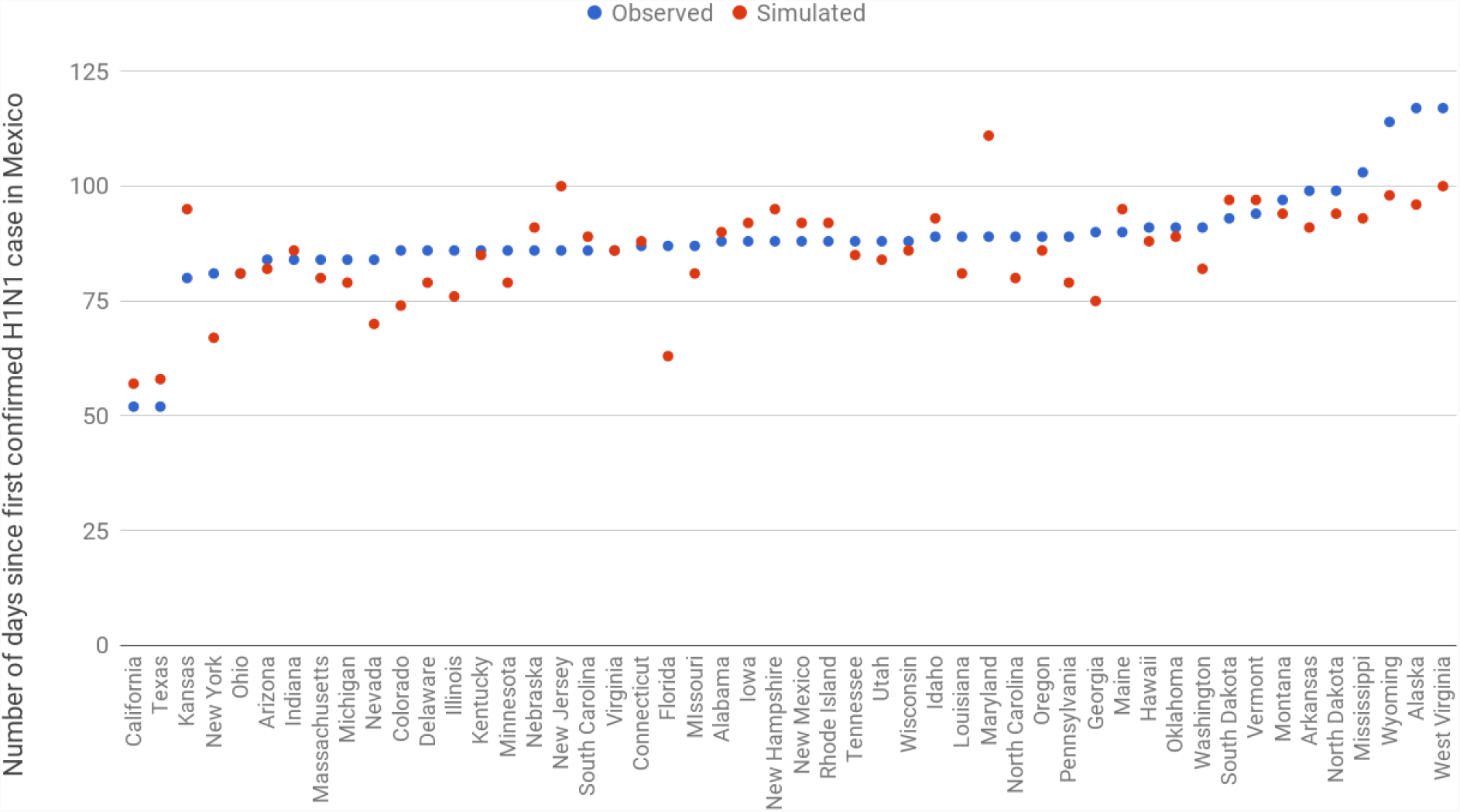
Observed^2^ and simulated date of H1N1 introduction into each State in the U.S.

The figure shows the observed and simulated date of the introduction of H1N1 in each of the 50 U.S. states sorted by increasing observed dates. The simulated dates are the average over 1,000 simulations of the stochastic metapopulation epidemic model.

A similar analysis was conducted at the global level, to compare the cumulative number of infected countries over time between the simulation and reported data, and illustrated in Figure S7. The difference between the observed^3^ and simulated results is minimal; the difference between the cumulative number of infected countries at each date, averaged over the first 100 days, is less than one. The maximum discrepancy during the first 100 days occurs 90 days after the start of the outbreak, at which time the simulation predicts six less countries to be infected.

**Figure S7.**
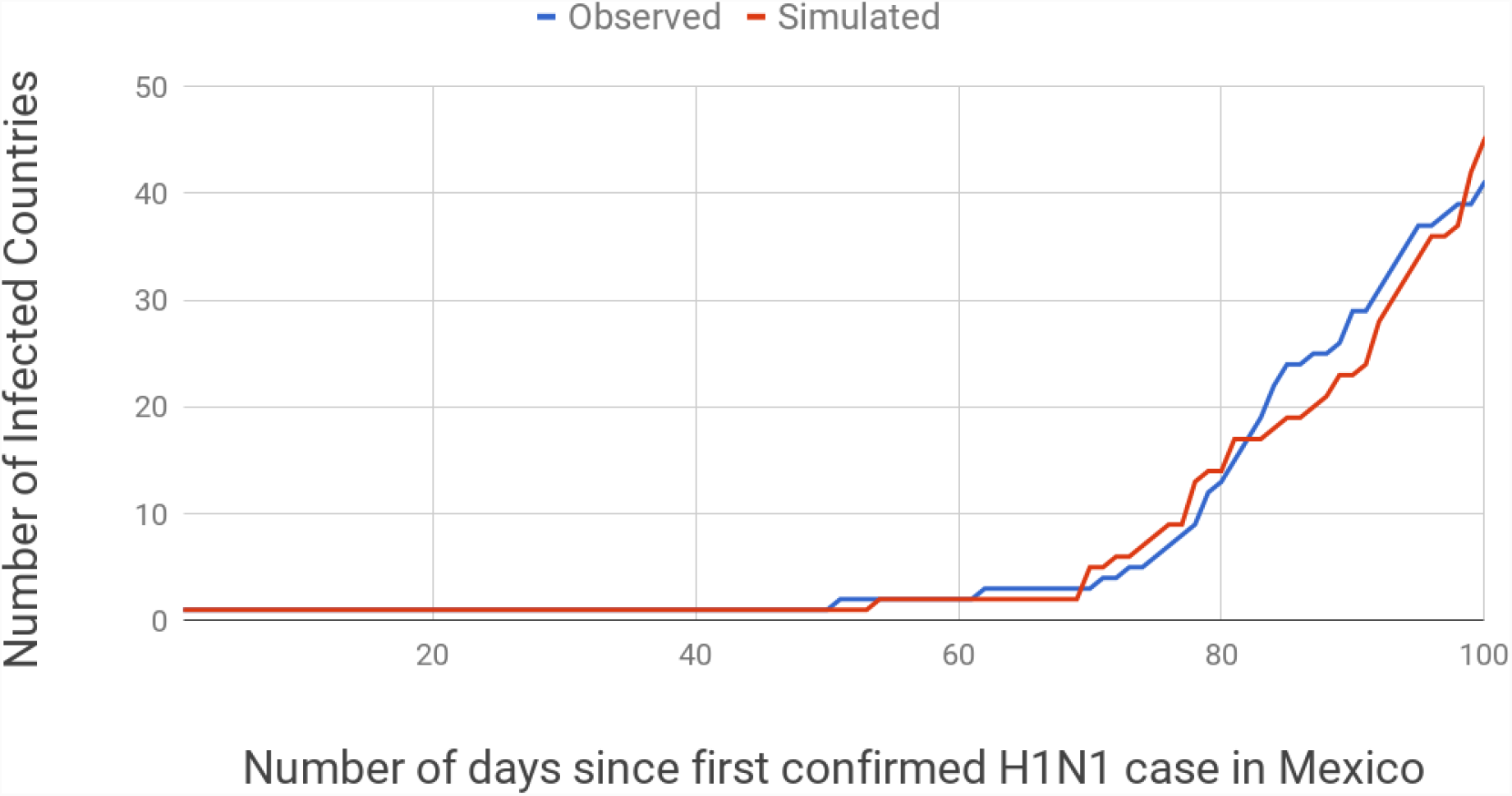
Observed and simulated cumulative number of countries infected over time. The observed curve is based on the date of arrival in country of first confirmed case**^3^** and the simulated curve is based on the expected date of first arrival averaged over 1,000 simulations of the stochastic metapopulation epidemic model.

To further validate the simulation model at a global level, explicit dates of first case arrivals for 10 countries known to be infected directly from Mexico are compared against the expected date of introduction from the simulation. Table S1^3,5^ highlights how closely the simulation captures the observed behavior of the outbreak, with an average difference of 5.4 days across the 10 countries. The observed and simulated arrival dates for all countries except Columbia fall within 8 days.

**Table S1.** Observed and simulated country infection dates ^3,5^. The simulated dates are the average of 1,000 simulations of the stochastic metapopulation epidemic model and rounded to the nearest date.

| Country | Observed<br>Arrival Date | Simulated<br>Arrival Date |
| --- | --- | --- |
| United States | 28/3/2009 | 30/3/2009 |
| United Kingdom | 21/4/2009 | 23/4/2009 |
| Australia | 9/5/2009 | 9/5/2009 |
| China | 2/5/2009 | 8/5/2009 |
| Canada | 8/4/2009 | 15/4/2009 |
| Spain | 22/4/2009 | 21/4/2009 |
| Germany | 28/4/2009 | 26/4/2009 |
| France | 1/5/2009 | 23/4/2009 |
| Colombia | 3/5/2009 | 15/4/2009 |
| Cuba | 25/4/2009 | 17/4/2009 |

### C. Airport control lists and frequency of control

To provide more details on the behavior of the proposed control strategies we analyse the sets of airports selected for control. For each control strategy, we report the list and the control level of all airports controlled in the base case scenarios MCO, PDX and HNL (note, airports are ordered by decreasing travel volume). We also report the frequency of choosing an airport among the 6 control strategies considered (all airports not listed in the tables are never selected for control and have thus a null frequency). We find that few airports are selected in all 6 control strategies. Even though the strategies have different metrics for ranking airports, EP and MC identify the same lists of airports for control, which explains their similar performance. The two lowest performing control strategies in terms of case numbers, LP and MT, have the most and least number of airports controlled, respectively. For LP, many controlled airports appear to not help in mitigating the spread of the disease, and for MT, the largest airports are always selected for control, which are also the most expensive, and therefore quickly deplete the budget available.

**Table S2a.** Airport control lists and frequency of airport control across control strategies in the MCO scenario.

| EP |  | MC |  | 1C |  | 1OU |  | LP |  | MT |  | Frequency |
| --- | --- | --- | --- | --- | --- | --- | --- | --- | --- | --- | --- | --- |
| List | Control | List | Control | List | Control | List | Control | List | Control | MT | Control |  |
| ATL | 1.00 | ATL | 1.00 | ATL | 1.00 | ATL | 1.00 | LAX | 1.00 | ATL | 1.00 | 6 |
| ORD | 1.00 | ORD | 1.00 | LAX | 0.38 | LAX | 1.00 | ORD | 1.00 | LAX | 1.00 | 5 |
| DFW | 1.00 | DFW | 1.00 | ORD | 1.00 | ORD | 1.00 | DFW | 1.00 | ORD | 1.00 | 4 |
| JFK | 1.00 | JFK | 1.00 | DFW | 1.00 | DFW | 1.00 | JFK | 1.00 | DFW | 1.00 | 3 |
| DEN | 1.00 | DEN | 1.00 | JFK | 1.00 | JFK | 1.00 | SFO | 1.00 | JFK | 1.00 | 2 |
| CLT | 1.00 | CLT | 1.00 | DEN | 1.00 | DEN | 1.00 | IAH | 1.00 | DEN | 1.00 | 1 |
| IAH | 1.00 | IAH | 1.00 | CLT | 1.00 | CLT | 1.00 | MIA | 0.86 | SFO | 1.00 |  |
| MIA | 0.06 | MIA | 0.06 | SEA | 1.00 | IAH | 1.00 | EWR | 1.00 | LAS | 1.00 |  |
| EWR | 1.00 | EWR | 1.00 | EWR | 1.00 | EWR | 1.00 | BOS | 1.00 | PHX | 1.00 |  |
| MSP | 1.00 | MSP | 1.00 | MSP | 1.00 | BOS | 1.00 | PHL | 1.00 | CLT | 1.00 |  |
| BOS | 1.00 | BOS | 1.00 | BOS | 1.00 | DTW | 1.00 | LGA | 1.00 | IAH | 1.00 |  |
| DTW | 1.00 | DTW | 1.00 | PHL | 1.00 | PHL | 1.00 | BWI | 1.00 | MIA | 1.00 |  |
| PHL | 1.00 | PHL | 1.00 | LGA | 1.00 | LGA | 1.00 | MDW | 1.00 | SEA | 0.77 |  |
| LGA | 1.00 | LGA | 1.00 | BWI | 1.00 | BWI | 1.00 | DCA | 1.00 |  |  |  |
| BWI | 1.00 | BWI | 1.00 | MDW | 1.00 | MDW | 0.46 | IAD | 1.00 |  |  |  |
| MDW | 1.00 | MDW | 1.00 | STL | 1.00 | SJU | 1.00 | SAN | 1.00 |  |  |  |
| DCA | 1.00 | DCA | 1.00 | SJU | 1.00 |  |  | DAL | 1.00 |  |  |  |
| SJU | 1.00 | SJU | 1.00 | IND | 1.00 |  |  | HOU | 1.00 |  |  |  |
|  |  |  |  | BDL | 1.00 |  |  | OAK | 1.00 |  |  |  |
|  |  |  |  | RHI | 1.00 |  |  | SNA | 1.00 |  |  |  |
|  |  |  |  | PAH | 1.00 |  |  | SJC | 1.00 |  |  |  |
|  |  |  |  | HIB | 1.00 |  |  | ONT | 1.00 |  |  |  |
|  |  |  |  | PKB | 1.00 |  |  | BUR | 1.00 |  |  |  |
|  |  |  |  |  |  |  |  | LGB | 1.00 |  |  |  |
|  |  |  |  |  |  |  |  | HPN | 1.00 |  |  |  |
|  |  |  |  |  |  |  |  | TTN | 1.00 |  |  |  |
|  |  |  |  |  |  |  |  | ILG | 1.00 |  |  |  |
|  |  |  |  |  |  |  |  | CLD | 1.00 |  |  |  |

**Table S2b.** Airport control lists and frequency of airport control across control strategies in the PDX scenario.

| EP |  | MC |  | 1C |  | 1OU |  | LP |  | MT |  | Frequency |
| --- | --- | --- | --- | --- | --- | --- | --- | --- | --- | --- | --- | --- |
| List | Control | List | Control | List | Control | List | Control | List | Control | MT | Control |  |
| ATL | 1.00 | ATL | 1.00 | ATL | 1.00 | ATL | 1.00 | LAX | 1.00 | ATL | 1.00 | 6 |
| LAX | 1.00 | LAX | 1.00 | LAX | 1.00 | LAX | 1.00 | ORD | 1.00 | LAX | 1.00 | 5 |
| ORD | 1.00 | ORD | 1.00 | ORD | 1.00 | ORD | 1.00 | DFW | 1.00 | ORD | 1.00 | 4 |
| DFW | 1.00 | DFW | 1.00 | DFW | 1.00 | DFW | 1.00 | JFK | 1.00 | DFW | 1.00 | 3 |
| JFK | 0.45 | JFK | 0.45 | JFK | 1.00 | JFK | 0.77 | SFO | 1.00 | JFK | 1.00 | 2 |
| DEN | 1.00 | DEN | 1.00 | DEN | 1.00 | DEN | 1.00 | IAH | 1.00 | DEN | 1.00 | 1 |
| SFO | 1.00 | SFO | 1.00 | SFO | 1.00 | SFO | 1.00 | MIA | 0.86 | SFO | 1.00 |  |
| LAS | 1.00 | LAS | 1.00 | LAS | 1.00 | LAS | 1.00 | EWR | 1.00 | LAS | 1.00 |  |
| PHX | 1.00 | PHX | 1.00 | PHX | 1.00 | PHX | 1.00 | BOS | 1.00 | PHX | 1.00 |  |
| SEA | 1.00 | SEA | 1.00 | SEA | 1.00 | SEA | 1.00 | PHL | 1.00 | CLT | 1.00 |  |
| MSP | 1.00 | MSP | 1.00 | MSP | 1.00 | MSP | 1.00 | LGA | 1.00 | IAH | 1.00 |  |
| SLC | 1.00 | SLC | 1.00 | SLC | 1.00 | SLC | 1.00 | BWI | 1.00 | MIA | 1.00 |  |
| SAN | 1.00 | SAN | 1.00 | HNL | 1.00 | SAN | 1.00 | MDW | 1.00 | SEA | 0.77 |  |
| HNL | 1.00 | HNL | 1.00 | OAK | 1.00 | HNL | 1.00 | DCA | 1.00 |  |  |  |
| OAK | 1.00 | OAK | 1.00 | SJC | 1.00 | OAK | 1.00 | IAD | 1.00 |  |  |  |
| SNA | 1.00 | SNA | 1.00 | SMF | 1.00 | SJC | 1.00 | SAN | 1.00 |  |  |  |
| SJC | 1.00 | SJC | 1.00 | GEG | 1.00 | SMF | 1.00 | DAL | 1.00 |  |  |  |
| SMF | 1.00 | SMF | 1.00 | DBQ | 1.00 |  |  | HOU | 1.00 |  |  |  |
| GEG | 1.00 | GEG | 1.00 | JLN | 1.00 |  |  | OAK | 1.00 |  |  |  |
| BOI | 1.00 | BOI | 1.00 | PAH | 1.00 |  |  | SNA | 1.00 |  |  |  |
|  |  |  |  | EKO | 1.00 |  |  | SJC | 1.00 |  |  |  |
|  |  |  |  | BRD | 1.00 |  |  | ONT | 1.00 |  |  |  |
|  |  |  |  | HYS | 1.00 |  |  | BUR | 1.00 |  |  |  |
|  |  |  |  |  |  |  |  | LGB | 1.00 |  |  |  |
|  |  |  |  |  |  |  |  | HPN | 1.00 |  |  |  |
|  |  |  |  |  |  |  |  | TTN | 1.00 |  |  |  |
|  |  |  |  |  |  |  |  | ILG | 1.00 |  |  |  |
|  |  |  |  |  |  |  |  | CLD | 1.00 |  |  |  |

**Table S2c.** Airport control lists and frequency of airport control across control strategies in the HNL scenario.

| EP |  | MC |  | 1C |  | 1OU |  | LP |  | MT |  | Frequency |
| --- | --- | --- | --- | --- | --- | --- | --- | --- | --- | --- | --- | --- |
| List | Control | List | Control | List | Control | List | Control | List | Control | MT | Control |  |
| ATL | 1.00 | ATL | 1.00 | ATL | 1.00 | ATL | 1.00 | LAX | 1.00 | ATL | 1.00 | 6 |
| LAX | 1.00 | LAX | 1.00 | LAX | 1.00 | LAX | 1.00 | ORD | 1.00 | LAX | 1.00 | 5 |
| ORD | 1.00 | ORD | 1.00 | ORD | 1.00 | ORD | 1.00 | DFW | 1.00 | ORD | 1.00 | 4 |
| DFW | 1.00 | DFW | 1.00 | DFW | 1.00 | DFW | 1.00 | JFK | 1.00 | DFW | 1.00 | 3 |
| JFK | 0.35 | JFK | 0.35 | JFK | 0.73 | JFK | 0.36 | SFO | 1.00 | JFK | 1.00 | 2 |
| DEN | 1.00 | DEN | 1.00 | DEN | 1.00 | DEN | 1.00 | IAH | 1.00 | DEN | 1.00 | 1 |
| SFO | 1.00 | SFO | 1.00 | SFO | 1.00 | SFO | 1.00 | MIA | 0.86 | SFO | 1.00 |  |
| LAS | 1.00 | LAS | 1.00 | LAS | 1.00 | LAS | 1.00 | EWR | 1.00 | LAS | 1.00 |  |
| PHX | 1.00 | PHX | 1.00 | PHX | 1.00 | PHX | 1.00 | BOS | 1.00 | PHX | 1.00 |  |
| IAH | 1.00 | IAH | 1.00 | IAH | 1.00 | SEA | 1.00 | PHL | 1.00 | CLT | 1.00 |  |
| SEA | 1.00 | SEA | 1.00 | SEA | 1.00 | MSP | 1.00 | LGA | 1.00 | IAH | 1.00 |  |
| SLC | 1.00 | SLC | 1.00 | IAD | 1.00 | SLC | 1.00 | BWI | 1.00 | MIA | 1.00 |  |
| SAN | 1.00 | SAN | 1.00 | SAN | 1.00 | SAN | 1.00 | MDW | 1.00 | SEA | 0.77 |  |
| PDX | 1.00 | PDX | 1.00 | PDX | 1.00 | HNL | 1.00 | DCA | 1.00 |  |  |  |
| OAK | 1.00 | OAK | 1.00 | SJC | 1.00 | PDX | 1.00 | IAD | 1.00 |  |  |  |
| SJC | 1.00 | SJC | 1.00 | OGG | 1.00 | OAK | 1.00 | SAN | 1.00 |  |  |  |
| SMF | 1.00 | SMF | 1.00 | KOA | 1.00 | SJC | 1.00 | DAL | 1.00 |  |  |  |
| OGG | 1.00 | OGG | 1.00 | LIH | 1.00 | OGG | 1.00 | HOU | 1.00 |  |  |  |
| GUM | 1.00 | GUM | 1.00 | ITO | 1.00 | KOA | 1.00 | OAK | 1.00 |  |  |  |
| KOA | 1.00 | KOA | 1.00 | ABE | 1.00 | LIH | 1.00 | SNA | 1.00 |  |  |  |
| LIH | 1.00 | LIH | 1.00 | LAN | 1.00 | ITO | 1.00 | SJC | 1.00 |  |  |  |
| ITO | 1.00 | ITO | 1.00 | SCE | 1.00 |  |  | ONT | 1.00 |  |  |  |
|  |  |  |  | LAW | 1.00 |  |  | BUR | 1.00 |  |  |  |
|  |  |  |  | ROW | 1.00 |  |  | LGB | 1.00 |  |  |  |
|  |  |  |  |  |  |  |  | HPN | 1.00 |  |  |  |
|  |  |  |  |  |  |  |  | TTN | 1.00 |  |  |  |
|  |  |  |  |  |  |  |  | ILG | 1.00 |  |  |  |
|  |  |  |  |  |  |  |  | CLD | 1.00 |  |  |  |

